# Binary and analog variation of synapses between cortical pyramidal neurons

**DOI:** 10.1101/2019.12.29.890319

**Authors:** Sven Dorkenwald, Nicholas L. Turner, Thomas Macrina, Kisuk Lee, Ran Lu, Jingpeng Wu, Agnes L. Bodor, Adam A. Bleckert, Derrick Brittain, Nico Kemnitz, William M. Silversmith, Dodam Ih, Jonathan Zung, Aleksandar Zlateski, Ignacio Tartavull, Szi-Chieh Yu, Sergiy Popovych, William Wong, Manuel Castro, Chris S. Jordan, Alyssa M. Wilson, Emmanouil Froudarakis, JoAnn Buchanan, Marc Takeno, Russel Torres, Gayathri Mahalingam, Forrest Collman, Casey Schneider-Mizell, Daniel J. Bumbarger, Yang Li, Lynne Becker, Shelby Suckow, Jacob Reimer, Andreas S. Tolias, Nuno Maçarico da Costa, R. Clay Reid, H. Sebastian Seung

**Affiliations:** Computer Science Department, Princeton University, Princeton, NJ, USA; Princeton Neuroscience Institute, Princeton University, Princeton, NJ, USA; Brain & Cognitive Sciences Department, Massachusetts Institute of Technology, Cambridge, MA, USA; Allen Institute for Brain Science, Seattle, WA, USA; Department of Neuroscience, Baylor College of Medicine, Houston, TX, USA; Center for Neuroscience and Artificial Intelligence, Baylor College of Medicine, Houston, TX, USA; Department of Electrical and Computer Engineering, Rice University, Houston, TX, USA

## Abstract

Learning from experience depends at least in part on changes in neuronal connections. We present the largest map of connectivity to date between cortical neurons of a defined type (L2/3 pyramidal cells), which was enabled by automated analysis of serial section electron microscopy images with improved handling of image defects. We used the map to identify constraints on the learning algorithms employed by the cortex. Previous cortical studies modeled a continuum of synapse sizes (Arellano et al. 2007) by a log-normal distribution (Loewenstein, Kuras, and Rumpel 2011; de Vivo et al. 2017; Santuy et al. 2018). A continuum is consistent with most neural network models of learning, in which synaptic strength is a continuously graded analog variable. Here we show that synapse size, when restricted to synapses between L2/3 pyramidal cells, is well-modeled by the sum of a binary variable and an analog variable drawn from a log-normal distribution. Two synapses sharing the same presynaptic and postsynaptic cells are known to be correlated in size (Sorra and Harris 1993; Koester and Johnston 2005; Bartol et al. 2015; Kasthuri et al. 2015; Dvorkin and Ziv 2016; Bloss et al. 2018; Motta et al. 2019). We show that the binary variables of the two synapses are highly correlated, while the analog variables are not. Binary variation could be the outcome of a Hebbian or other synaptic plasticity rule depending on activity signals that are relatively uniform across neuronal arbors, while analog variation may be dominated by other influences. We discuss the implications for the stability-plasticity dilemma.

## Introduction

Synapses between excitatory neurons in the cortex and hippocampus are typically made onto spines, tiny thorn-like protrusions from dendrites (Yuste 2010). In the 2000s, long-term *in vivo* microscopy studies showed that dendritic spines change in shape and size, and can appear and disappear (Holtmaat and Svoboda 2009; Bhatt, Zhang, and Gan 2009). Spine dynamics were interpreted as synaptic plasticity, because spine volume is well-correlated with physiological strength of a synapse (Matsuzaki et al. 2001; Noguchi et al. 2011; Holler et al. 2021). The plasticity was thought to be in part activity-dependent, because spine volume increases with long-term potentiation (Matsuzaki et al. 2004; Kopec et al. 2006; Noguchi et al. 2019). Given that the sizes of other synaptic structures (postsynaptic density, presynaptic active zone, and so on) are well-correlated with spine volume and with each other (Harris and Stevens 1989), we use the catch-all term “synapse size” to refer to the size of any synaptic structure, and “synapse strength” as a synonym.

In the 2000s, some hypothesized that long-term plasticity involves discrete transitions of synapses between two structural states (Kasai et al. 2003; Bourne and Harris 2007). Quantitative studies of cortical synapses, however, found no evidence for discreteness (Arellano et al. 2007; Loewenstein, Kuras, and Rumpel 2011; Loewenstein, Yanover, and Rumpel 2015; de Vivo et al. 2017; Santuy et al. 2018; Harris and Stevens 1989). Whether in theoretical neuroscience or artificial intelligence, it is common for the synaptic strengths in a neural network model to be continuously variable, enabling learning to proceed by the accumulation of arbitrarily small synaptic changes over time.

Here we reexamine the discrete versus continuous dichotomy using a wiring diagram between 334 layer 2/3 pyramidal cells (L2/3 PyCs) reconstructed from serial section electron microscopy (ssEM) images of mouse primary visual cortex. We show that synapses between L2/3 PyCs are well-modeled as a binary mixture of log-normal distributions. If we further restrict consideration to dual connections, two synapses sharing the same presynaptic and postsynaptic cells, the binary mixture exhibits a statistically significant bimodality. It is therefore plausible that the binary mixture reflects two underlying structural states, and is more than merely an improvement in curve fitting.

According to our best fitting mixture model, synapse size is the sum of a binary variable and a log-normal continuous variable. To probe whether these variables are modified by synaptic plasticity, we again examine dual connections. It was previously shown that synapse pairs at dual connections are correlated in size, and the correlations have been attributed to activity-dependent plasticity (Sorra and Harris 1993; Koester and Johnston 2005; Bartol et al. 2015; Kasthuri et al. 2015; Dvorkin and Ziv 2016; Bloss et al. 2018; Motta et al. 2019). We find that the binary variables are highly correlated, while the continuous variables are not. This suggests that binary variation is the outcome of a Hebbian or other synaptic plasticity rule driven by activity signals that are relatively uniform across neuronal arbors, while analog variation is dominated by other factors.

Our findings require the specificity of our synaptic population. If we expand the analysis to include a broader population of cortical synapses, bimodality is no longer observed. This likely explains why previous studies have supported the continuum idea of cortical synapses (Arellano et al. 2007; Loewenstein, Kuras, and Rumpel 2011; de Vivo et al. 2017; Santuy et al. 2018; Ofer et al. 2021; Harris and Stevens 1989).

The specificity of our synaptic population was made possible because each of the 334 neurons taking part in the 1735 connections in our cortical wiring diagram could be identified as a L2/3 PyC based on a soma and sufficient dendrite and axon contained in the ssEM volume. The closest precedents for wiring diagrams between cortical neurons of a defined type had 29 connections between 43 L2/3 PyCs in mouse visual cortex (W.-C. A. Lee et al. 2016), 63 connections between 22 L2 excitatory neurons in mouse medial entorhinal cortex (Schmidt et al. 2017), and 32 connections between 89 L4 neurons in mouse somatosensory cortex (Motta et al. 2019).

Our cortical reconstruction is publicly available (https://www.microns-explorer.org/phase1)(Turner, Macrina, et al. 2020; Schneider-Mizell et al. 2020) and the code that generated the reconstruction is already freely available (Data and Code Availability).

## Handling of ssEM image defects

We acquired a 250×140×90 μm^3^ ssEM dataset (Extended Data Fig. 1) from L2/3 primary visual cortex of a P36 male mouse. When we aligned a pilot subvolume and applied state-of-the-art convolutional nets, we found many reconstruction errors, mainly due to misaligned images and damaged or incompletely imaged sections. This was disappointing given reports that convolutional nets can approach human-level performance on one benchmark ssEM image dataset (Beier et al. 2017; Zeng, Wu, and Ji 2017). The high error rate could be explained by the fact that image defects are difficult to escape in large volumes, though they may be rare in small (<1000 μm^3^) benchmark datasets.

Indeed, ssEM images were historically considered problematic for automated analysis (Briggman and Bock 2012; K. Lee et al. 2019) because they were difficult to align, contained defects caused by lost or damaged serial sections, and had inferior axial resolution (Knott et al. 2008). These difficulties were the motivation for developing block face electron microscopy (bfEM) as an alternative to ssEM (Denk and Horstmann 2004). Most large-scale ssEM reconstructions have been completely manual, while many large-scale bfEM reconstructions have been semiautomated (19/20 and 5/10 in Table 1 of (Kornfeld and Denk 2018)). On the other hand, the higher imaging throughput of ssEM (Nickell and Zeidler 2019; Yin et al. 2019) makes it suitable for scaling up to volumes that are large enough to encompass the arbors of mammalian neurons.

We supplemented existing algorithms for aligning ssEM images (Saalfeld et al. 2012) with human-in-the-loop capabilities. After manual intervention by a human expert, large misalignments were resolved but small ones still remained near damaged locations and near the borders of the volume. Therefore we augmented the training data for our convolutional net with simulated misalignments and missing sections (Fig. 1a, Extended Data Fig. 2). The resulting net was better able to trace neurites through such image defects (Fig. 1b, quantification in Supplementary Information 1). Other methods for handling ssEM image defects are being proposed (Li et al. 2019), and we can look forward to further gains in automated reconstruction accuracy in the future.

**Figure 1:**
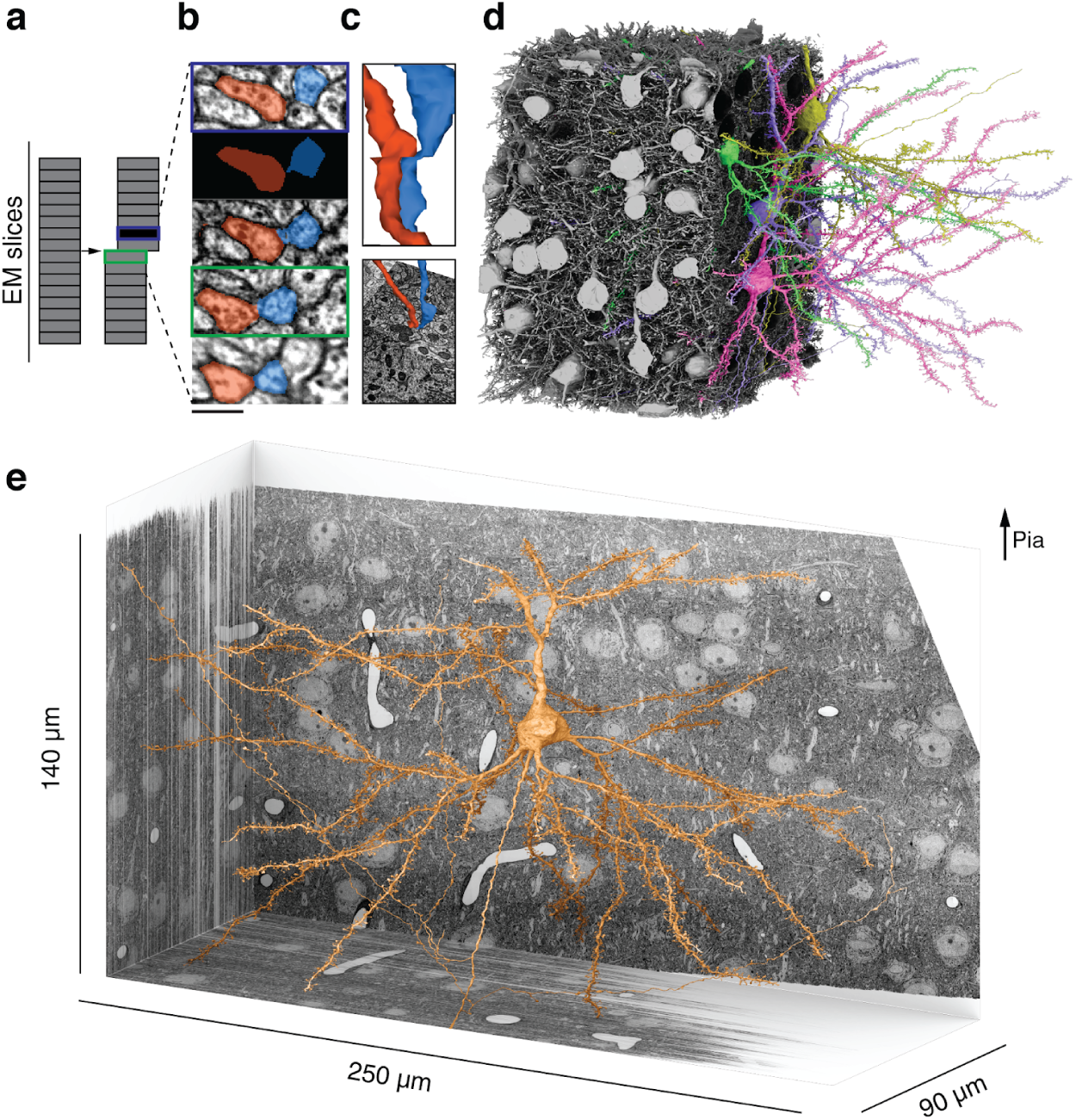
Reconstructing cortical circuits in spite of ssEM image defects. **(a)** Ideally, imaging serial sections followed by computational alignment would create an image stack that reflects the original state of the tissue (left). In practice, image stacks end up with missing sections (blue) and misalignments (green). Both kinds of defects are easily simulated when training a convolutional net to detect neuronal boundaries. Small subvolumes are depicted rather than the entire stack, and image defects are typically local rather than extending over an entire section. **(b)** The resulting net can trace more accurately, even in images not previously seen during training. Here a series of five sections contains a missing section (blue frame) and a misalignment (green). The net “imagines” the neurites through the missing section, and traces correctly in spite of the misalignment. **(c)** 3D reconstructions of the neurites exhibit discontinuities at the misalignment, but are correctly traced. **(d)** All 362 pyramidal cells with somas in the volume (gray), cut away to reveal a few examples (colors). **(e)** L2/3 pyramidal cell reconstructed from ssEM images of mouse visual cortex. Scale bars: 300 nm (b)

## Wiring diagram between L2/3 pyramidal cells

After alignment and automatic segmentation (Methods), we semi-automatically identified 417 pyramidal cells (PyCs) with somas in the volume based on morphological characteristics and automated nucleus detection and automated nucleus detection (Fig. 1d, e, Methods). We then chose a subset of 362 PyCs with sufficient neurite length within the volume for proofreading. Remaining errors in the segmentation of these PyCs were corrected using an interactive system that enabled human experts to split and merge objects.

Overall, the PyC reconstructions were corrected through ∼1,300 hours of human proofreading to yield 670 mm cable length (axon: 100 mm, dendrite: 520 mm, perisomatic: 40 mm, Extended Data Fig. 2). We examined 12 randomly sampled axons and conservatively estimated 0.28 merge errors per millimeter remain after proofreading (see Methods for other estimates). The dendrites of the PyCs receive more than one quarter of the 3.2 million synapses that were automatically detected in the volume (Methods, (Turner, Lee, et al. 2020)). However, the synapses onto PyC dendrites are almost all from “orphan” axons, defined as those axonal fragments that belong to somas of unknown location outside the volume. Using these automatically detected synapses as a starting point, we mapped all connections between this set (Methods). The end result was a wiring diagram of 1,960 synapses from 1,735 connections between 334 L2/3 PyCs in the dataset (Fig. 2a). Note that some connections are multisynaptic, i.e., they are mediated by multiple synapses sharing the same presynaptic and postsynaptic cells (Fig. 2b, Extended Data Fig. 3, see Supplementary Information 7 for a tabular overview of these statistics).

**Figure 2:**
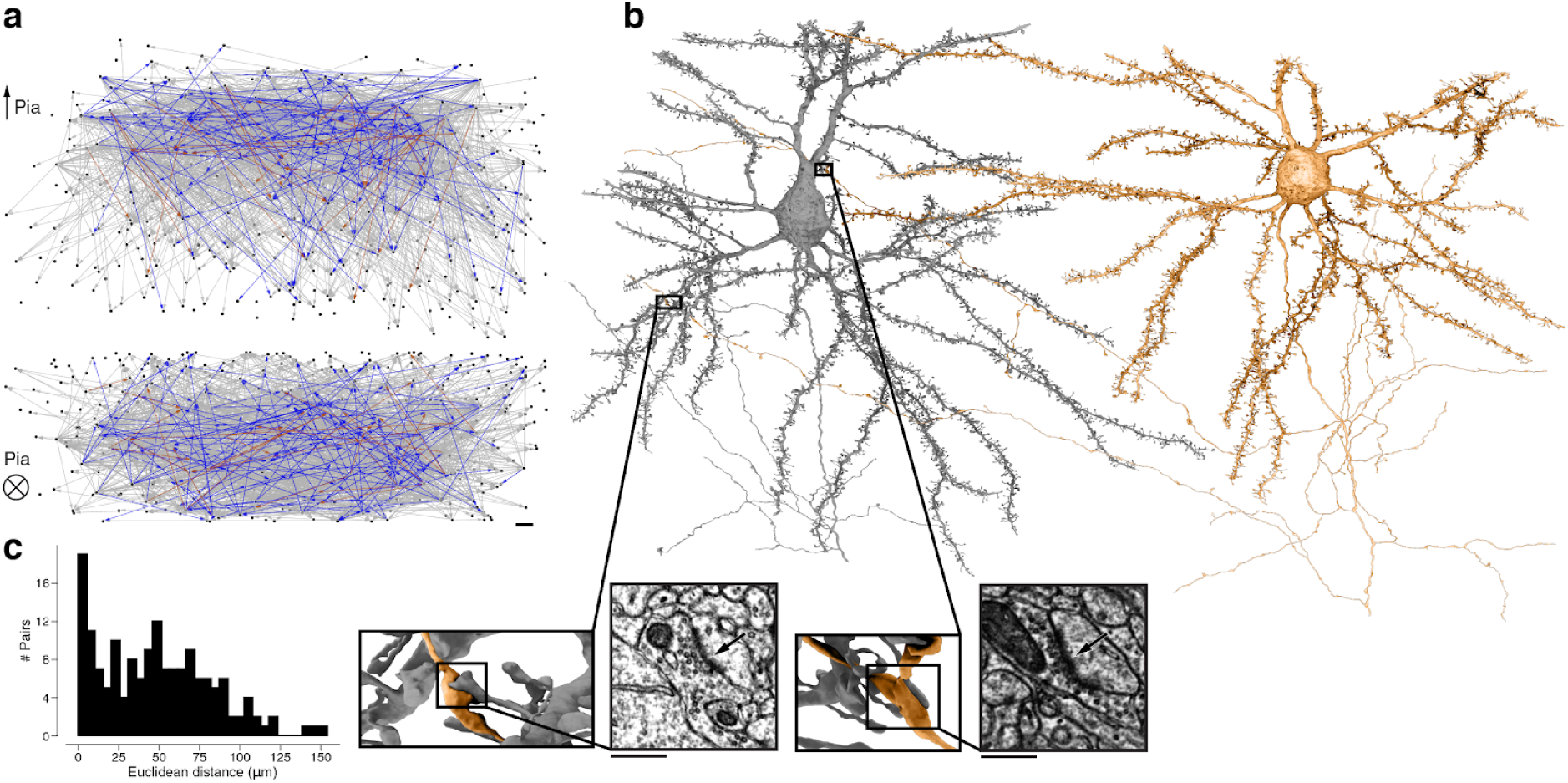
Wiring diagram for cortical neurons including multisynaptic connections. **(a)** Wiring diagram of 362 proofread L2/3 pyramidal cells as a directed graph. Two orthogonal views with nodes at 3D locations of cell bodies. Single (gray), dual (blue), and triple, quadruple, quintuple (red) connections. **(b)** Dual connection from a presynaptic cell (orange) to a postsynaptic cell (gray). Ultrastructure of both synapses can be seen in closeups from the EM images. The euclidean distance between the synapses is 64.3 μm. **(c)** Distribution of Euclidean distances between synapse pairs of dual connections. Median distance is 46.5 μm. Scale bars: 10 μm (a), 500 nm (b)

For clarity, we emphasize that our usage of the term “multisynaptic” refers to multiple synapses in parallel rather than in series. A connection usually (89.1%) contains one synapse, but can contain up to five synapses (2: 9.22%, 3: 1.38%, 4: 0.17%, 5: 0.12%).^1^ The dimensions of our reconstruction allowed us to not only observe dual connections with synapses up to 10 μm apart (Bartol et al. 2015; Kasthuri et al. 2015; Bloss et al. 2018) but also dual connections with two synapses more than 100 μm apart (Fig. 2b,c) (W.-C. A. Lee et al. 2016).

## Binary latent states

Previous studies of cortical synapses have found a continuum of synapse sizes (Arellano et al. 2007) that is well-modeled by a log-normal distribution (Loewenstein, Kuras, and Rumpel 2011; de Vivo et al. 2017; Santuy et al. 2018). Even researchers who report bimodally distributed synapse size in hippocampus (Spano et al. 2019) still find log-normally distributed synapse size in neocortex (de Vivo et al. 2017) by the same methods.

We quantified the size of each synapse by the volume of the spine head (Fig. 2b, 3a)^2^. In the following, “spine volume” will serve as a synonym for spine head volume. The distribution of spine volumes is highly skewed, with a long tail of large spines (Fig. 3b) as observed before (Loewenstein, Kuras, and Rumpel 2011; Santuy et al. 2018). Because of the skew, it is helpful to visualize the distribution using a logarithmic scale for spine volume (Loewenstein, Kuras, and Rumpel 2011; Bartol et al. 2015). We were surprised to find that the distribution deviated from normality, due to a “knee” on the right side of the histogram (Fig. 3c). A mixture of two normal distributions was a better fit than a single normal distribution when accounting for the number of free parameters (likelihood ratio test: p<1e-39, n=1960, Methods).

**Figure 3:**
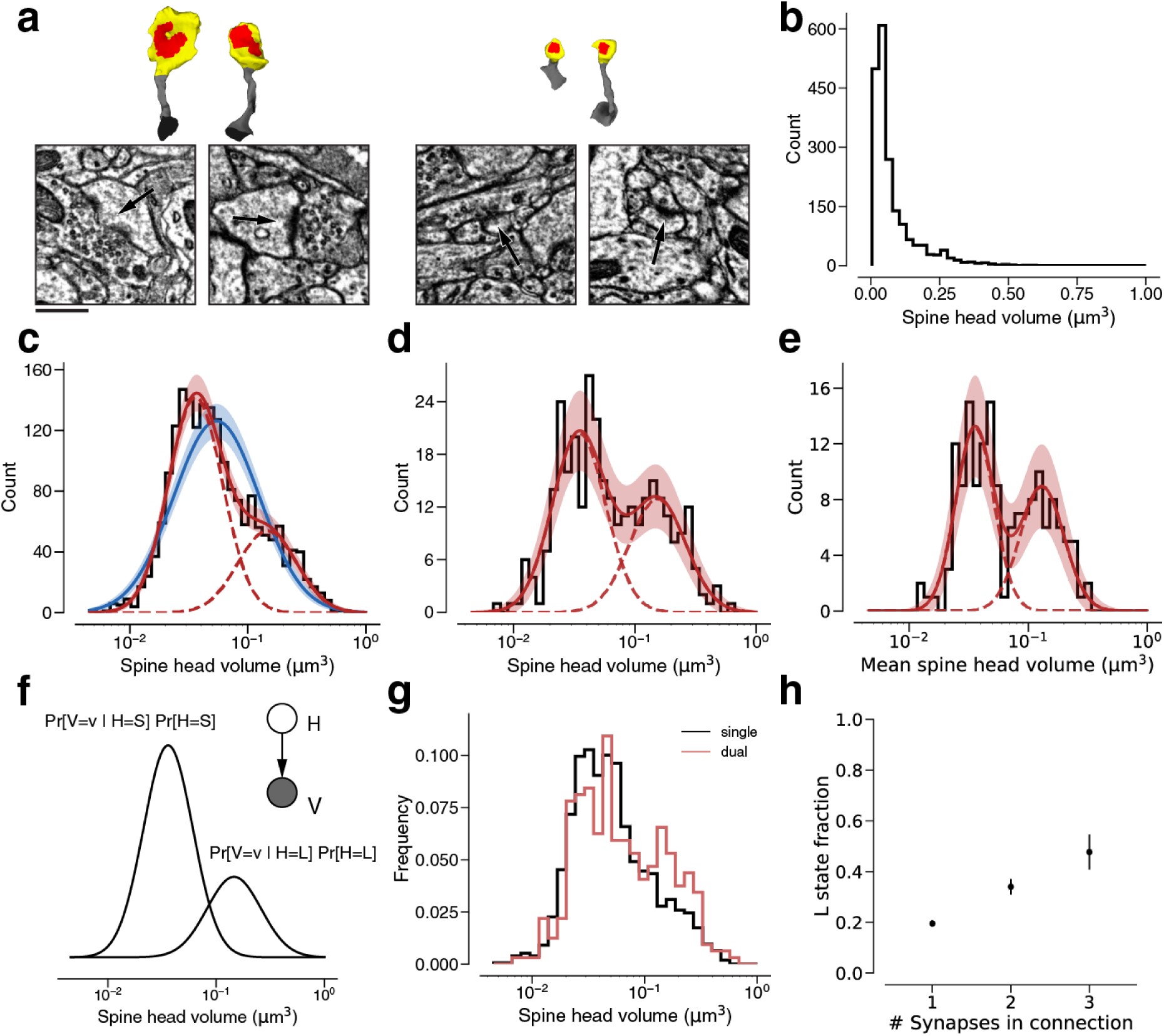
Modeling spine head volume with a mixture of two log-normal distributions. **(a)** Dendritic spine heads (yellow) and clefts (red) of dual connections between L2/3 PyCs. The associated EM cutout shows a 2D slice through the synapse. The synapses are centered in the EM images. **(b)** Skewed histogram of spine volume for all 1960 recurrent synapses between L2/3 PyCs, with a long tail of large spines. **(c)** Histogram of the spine volumes in (b), logarithmic scale. A mixture (red, solid) of two log-normal distributions (red, dashed) fits better (likelihood ratio test, p<1e-39, n=1960) than a single normal (blue). **(d)** Spine volumes belonging to dual connections between L2/3 PyCs, modeled by a mixture (red, solid) of two log-normal distributions (red, dashed). **(e)** Dual connections between L2/3 PyCs, each summarized by the geometric mean of two spine volumes, modeled by a mixture (red, solid) of two log-normal distributions (red, dashed). **(f)** Mixture of two normal distributions as a probabilistic latent variable model. Each synapse is described by a latent state *H* that takes on values “S” and “L” according to the toss of a biased coin. Spine volume *V* is drawn from a log-normal distribution with mean and variance determined by latent state. The curves shown here represent the best fit to the data in (d). Heights are scaled by the probability distribution of the biased coin, known as the mixture weights. **(g)** Comparison of spine volumes for single (black) and dual (red) connections. **(h)** Probability of the “L” state (mixture weight) versus number of synapses in the connection. Error bars are standard deviations estimated by bootstrap sampling. Scale bar: 500 nm (a). Error bars are ± 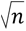 of the model fit (c, d, e) and standard deviation from bootstrapping (h).

We next restricted our consideration to the 320 synapses belonging to 160 dual connections between the PyCs. Again, a binary mixture of normal distributions was a better fit (Fig. 3d) than a single normal distribution (normal fit not shown, likelihood ratio test: p<1e-7, n=320). Next, we made use of the fact that synapses from dual connections are paired. For each pair, we computed the geometric mean (i.e. mean in log-space) of spine volumes and found that this quantity is also well-modeled by a binary mixture of normal distributions (Fig. 3e, see Supplementary Information 3 for the arithmetic mean, Supplementary Information 5 for histograms without model fits and Supplementary Information 8 for fit results).

A binary mixture model might merely be a convenient way of approximating deviations from normality. We would like to know whether the components of our binary mixture really have a biological basis, i.e., whether they correspond to two structural states of synapses. A mixture of two normal distributions can be unimodal or bimodal, depending on the model parameter^3^ (Robertson and Fryer 1969). When comparing best fit unimodal and bimodal mixtures we found that a bimodal model yields a significantly superior fit for spine volume and geometric mean of spine volume (Holzmann and Vollmer 2008) (Extended Data Fig. 4). This bimodality makes it plausible that the mixture components correspond to biological states of synapses. According to this interpretation, synapses are drawn from two latent states (Fig. 3f). In “S” and “L” states, spine volumes are drawn from log-normal distributions with small and large means, respectively. It should be noted that there is some overlap between mixture components (Fig. 3f), so that an S synapse can be larger than an L synapse.

To validate this finding, we quantified synaptic cleft size as the number of voxels labeled by the output of our automated cleft detector (Supplementary Information 4). We found a close relationship between spine volume and cleft size in our data (Extended Data Fig. 5a), in accord with previous studies (Harris and Stevens 1989; Arellano et al. 2007; Bartol et al. 2015). When spine volume is replaced by cleft size in the preceding analysis, we obtain similar results (Extended Data Fig. 5).

To probe possible model dependence on the number of synapses per connection, we separated our synapses into populations drawn from single, dual and triple connections, and fit an independent binary mixture model to each population. The parameters of the mixture components for dual and triple connections were not significantly different from the parameters for single connections (Supplementary Information 2). When comparing these distributions we observed an overrepresentation of large synapses for dual connections compared to single connections (Fig. 3g). We wondered if the previously reported mean spine volume increase with the number of synapses per connection (Supplementary Information 2, (Bloss et al. 2018) could be explained with a synapse redistribution between the latent states. This time, we only fit the component weights to single, dual, triple connections while keeping the Gaussian components constant (see Methods). We found a linear increase in fraction of synapses in the “L” state with the number of synapses per connections (Fig. 3h).^4^

Large spines have been reported to contain an intracellular organelle called a spine apparatus (SA), which is a specialized form of smooth endoplasmic reticulum (ER) (Peters and Kaiserman-Abramof 1970; Spacek 1985; Harris and Stevens 1989). We manually annotated SA in all dendritic spines of all synapses between L2/3 PyCs, and confirmed quantitatively that the probability of a spine apparatus increases with spine volume (Extended Data Fig. 6, Methods).

## Correlations at dual connections

Positive correlation between synapse sizes at dual connections has been reported previously in hippocampus (Sorra and Harris 1993; Bartol et al. 2015; Bloss et al. 2018) and neocortex (Kasthuri et al. 2015; Motta et al. 2019). According to our binary mixture model, synapse size is the sum of a binary variable and a log-normal continuous variable. We decided to quantify the contributions of these variables to synapse size correlations.

The dendritic spines for all dual connections between L2/3 PyCs are rendered in Extended Data Fig. 3. In a scatter plot of spine volumes (Fig. 4a, see Supplementary Information 6 for an unoccluded plot), positive correlations are evident (Pearson’s *r*=0.418). We fit the joint distribution of the spine volumes by a mixture model like Fig. 3f, while allowing the latent states to be correlated (Fig. 4a, f, see Supplementary Information 9 for fit results, Extended Data Fig. 7). In the best-fitting model, SS occurs roughly half the time, LL one third of the time, and the mixed states (SL, LS) occur more rarely (Fig. 4e). The low probability of the mixed states can be seen directly in the scarcity of points in the upper left and lower right corners of the scatter plot (Fig. 4a). Pearson’s phi coefficient, the specialization of Pearson’s correlation coefficient to binary variables, is 0.637.

**Figure 4:**
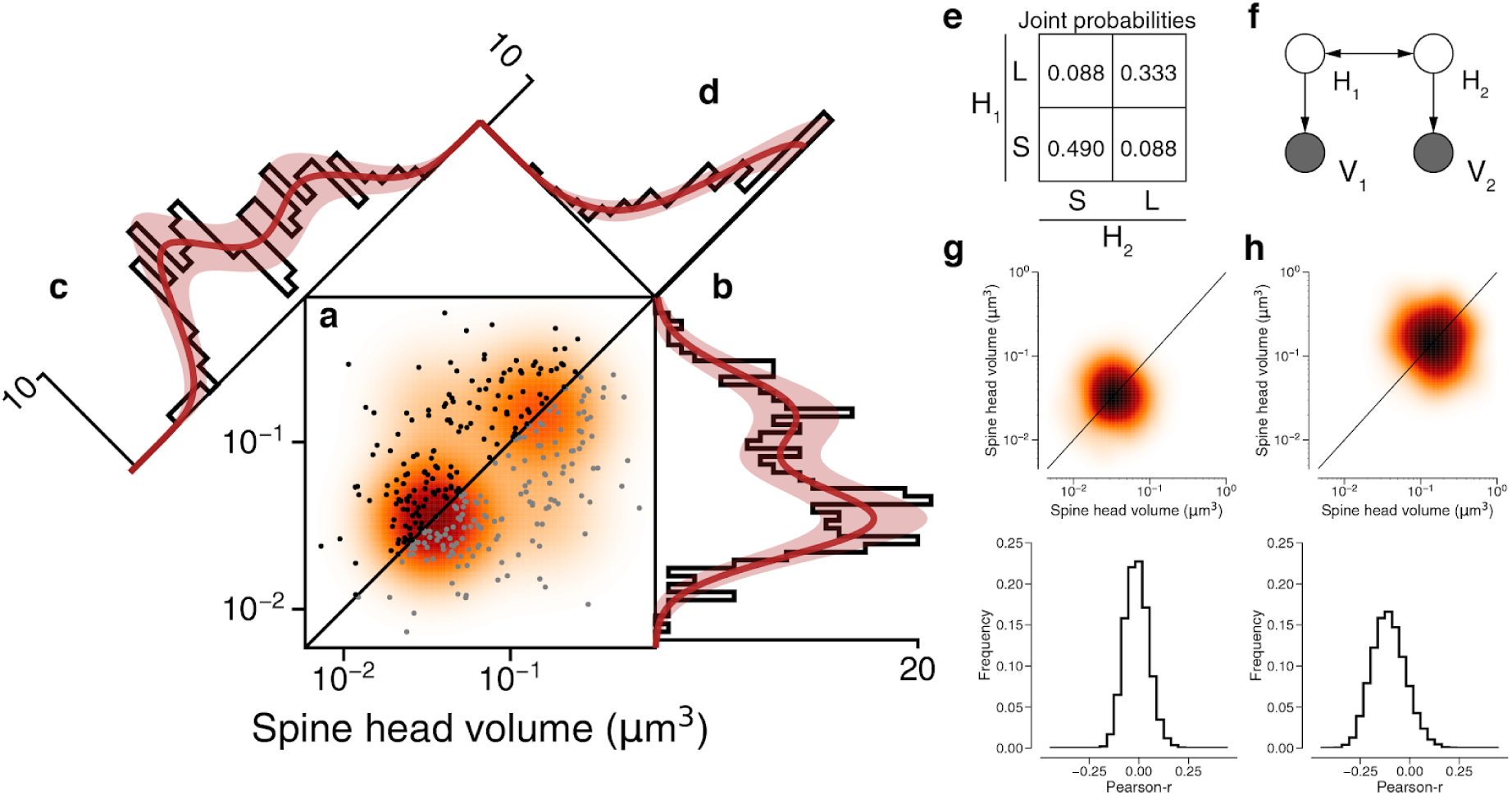
Latent state correlations between spines at dual connections. **(a)** Scatter plot of spine volumes (black, lexicographic ordering) for dual connections. Data points are mirrored across the diagonal (gray). The joint distribution is fit by a mixture model (orange) like that of Fig. 3f, but with latent states correlated as in (e). **(b)** Projecting the points onto the vertical axis yields a histogram of spine volumes for dual connections (cf. Fig. 3d). Model is derived from the joint distribution. **(c)** Projecting onto the x=y diagonal yields a histogram of the geometric mean of spine volumes (cf. Fig. 3e). Model is derived from the joint distribution. **(d)** Projecting onto the x=−y diagonal yields a histogram of the ratio of spine volumes. **(e)** The latent states of synapses in a dual connection (*H*_1_ and *H*_2_) are more likely to be the same (SS or LL) than different (SL/LS), as shown by the joint probability distribution. **(f)** When conditioned on the latent states, the spine volumes (*V*_1_ and *V*_2_) are statistically independent, as shown in this dependency diagram of the model. **(g)**, **(h)** Sampling synapse pairs to SS and LL states according to their state probabilities. The top shows a kernel density estimation of multiple iterations of sampling. The bottom shows the distribution of pearson-r correlations across many sampling rounds (N=10,000). Error bars are ± 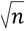 of the model fit.

Our mixture model assumes that the spine volumes are independent when conditioned on the latent states. To visualize whether this assumption is justified by the data, Fig. 4 shows 1D projections of the joint distribution onto different axes. The projection onto the vertical axis (Fig. 4b) is the marginal distribution, the overall size distribution for all synapses that belong to dual connections (same as Fig. 3d). The projection onto the *x*=*y* diagonal (Fig. 4c) is the distribution of the geometric mean of spine volume for each dual connection (same as Fig. 3e). The projection onto the *x*=−*y* diagonal (Fig. 4d) is the distribution of the ratio of spine volumes for each dual connection. For all three projections, the good fit suggests that the data are consistent with the mixture model’s assumption of isotropic normal distributions for the LL and SS states.^5^

For a quantitative test of the isotropy assumption, we resampled observed data points with weightings computed from the posterior probabilities of the SS and LL states (Fig. 4g, h). If the model were consistent with the data, the resampled data would obey an isotropic normal distribution. Indeed, Pearson’s correlation for the resampled data is not significantly different from zero (Fig. 4g, h). Therefore the spine volumes in a dual connection are approximately uncorrelated when conditioned on the latent states.

## Specificity of latent state correlations

Could the observed correlations between synapses in dual connections be caused by crosstalk between plasticity of neighboring synapses (<10 μm separation), which has been reported previously (Harvey and Svoboda 2007; Harvey et al. 2008)? We looked for dependence of latent state correlations on separation by splitting dual connections into two groups, those with synapses nearer or farther than the median Euclidean distance of 46.5 μm. Both groups were fit by mixture models with positive correlations between latent variables (near: *φ* =0.53, far: *φ* = 0.75, see Methods, Extended Data Fig. 8). In other words, for dual connections involving pairs of distant synapses, the latent state correlations are still strong.

We also considered the possibility of correlations in pairs of synapses sharing the same presynaptic cell but not the same postsynaptic cell, or pairs of synapses sharing the same postsynaptic cell but not the same presynaptic cell (Bartol et al. 2015; Kasthuri et al. 2015; Dvorkin and Ziv 2016; Bloss et al. 2018; Motta et al. 2019). We randomly drew such synapse pairs from the set of synapses that belong to dual connections (and hence belong to PyCs that participate in dual connections). Correlations in the latent state or synapse size were negligible (same axon: φ= −0.11 ± 0.08 SD, *r* = −0.06 ± 0.06 SD; same dendrite: φ= −0.06 ± 0.06 SD, *r* = −0.13 ± 0.05 SD; Extended Data Fig. 9), similar to previous findings (Bloss et al. 2018; Motta et al. 2019).

## Discussion

Our synapse size correlations are specific to pairs of synapses that share both the same presynaptic and postsynaptic L2/3 PyCs, analogous to previous findings (Sorra and Harris 1993; Koester and Johnston 2005; Bartol et al. 2015; Kasthuri et al. 2015; Dvorkin and Ziv 2016; Bloss et al. 2018; Motta et al. 2019). We have further demonstrated that the correlations exist even for large spatial separations between synapses. More importantly, we have shown the correlations are confined to the binary latent variables in our synapse size model; the log-normal analog variables exhibit little or no correlation.

The correlations in the binary variables could arise from a Hebbian or other synaptic plasticity rule driven by presynaptic and postsynaptic activity signals that are relatively uniform across neuronal arbors. Such signals are shared by synapses in a multisynaptic connection (Sorra and Harris 1993; Koester and Johnston 2005; Bartol et al. 2015; Kasthuri et al. 2015; Dvorkin and Ziv 2016; Bloss et al. 2018; Motta et al. 2019).

We speculate that much of the analog variation arises from the spontaneous dynamical fluctuations that have been observed at single dendritic spines through time-lapse imaging. Computational models of this temporal variance suggest that it can account for much of the population variance (Yasumatsu et al. 2008; Loewenstein, Kuras, and Rumpel 2011; Statman et al. 2014). Experiments have shown that large dynamical fluctuations persist even after activity is pharmacologically blocked (Yasumatsu et al. 2008; Statman et al. 2014). Another possibility is that the analog variation arises from plasticity driven by activity-related signals that are local to neighborhoods within neuronal arbors.

It has been argued that the observed structural volatility of synapses is challenging to reconcile with the stability of memory (Loewenstein, Kuras, and Rumpel 2011). Our findings suggest two possible resolutions of the stability-plasticity dilemma. In one scenario, synapses only appear volatile because fluctuations in the analog variable obscure the stability of the binary variable. This scenario is consistent with the idea that synapses behave like binary switches that are flipped by activity-dependent plasticity. Switch-like behavior could arise from bistable networks of molecular interactions at synapses (Lisman 1985), has been observed in physiology experiments on synaptic plasticity (Petersen et al. 1998; O’Connor, Wittenberg, and Wang 2005), and has been the basis of a number of computational models of memory (Tsodyks 1990; Amit and Fusi 1994; Fusi, Drew, and Abbott 2005).

In a second scenario, there is no need for synapses to be intrinsically stable as learning from experience is always ongoing in a sensory cortical area. Activity-dependent plasticity causes synapses to partition into two clusters located at upper and lower bounds for synaptic size (S. Song, Miller, and Abbott 2000; Van Rossum, Bi, and Turrigiano 2000; Rubin, Lee, and Sompolinsky 2001). In this scenario, synapses are intrinsically continuous and volatile, and the binary mixture is an outcome of ongoing learning.

Bimodality and strong correlations were found for a restricted ensemble of synapses, those belonging to dual connections between L2/3 PyCs. However, bimodality is not observed for the ensemble of all excitatory synapses onto L2/3 PyCs, including those from orphan axons (Extended Data Fig. 10). This ensemble is similar to ones studied previously, i.e., synapses onto L2/3 PyCs (Arellano et al. 2007), L4 neurons (Motta et al. 2019) or L5 PyCs (Loewenstein, Kuras, and Rumpel 2011). Bimodality and strong correlations are also not observed for the ensemble of all dual connections received by L2/3 PyCs, including those from orphan axons (Extended Data Fig. 10). Because our novel findings are based on a highly specific population of synapses, they are not inconsistent with previous studies that failed to find evidence for discreteness of cortical synapses (Arellano et al. 2007; Loewenstein, Kuras, and Rumpel 2011; Loewenstein, Yanover, and Rumpel 2015; de Vivo et al. 2017; Santuy et al. 2018).

Bimodality and correlations may turn out to be heterogeneous across classes of neocortical synapses. Heterogeneity in the hippocampus has been demonstrated by the finding that dual connections onto granule cell dendrites in the middle molecular layer of dentate gyrus (Bromer et al. 2018) do not exhibit the correlations that are found in stratum radiatum of CA1 (Bartol et al. 2015; Bloss et al. 2018).

Since the physiological strength of a multisynaptic connection can be approximately predicted from the sum of synaptic sizes (Holler-Rickauer, Koestinger, and Martin 2019), our S and L latent states and their correlations have implications for the debate over whether infrequent strong connections play a disproportionate role in cortical computation (Sen Song et al. 2005; Cossell et al. 2015; Scholl et al. 2019).

Testing the generality of our findings would be facilitated by further scale-up of connectomics, which in turn would be aided by additional innovations in handling of ssEM image defects. Our experiences with ssEM data are paralleled by other efforts to transform artificial intelligence research into real-world systems. For example, consider that self-driving cars may perform impressively under “normal” conditions, but fail in the so-called “edge” or “corner” cases (Madrigal 2018) that currently stand in the way of fully autonomous operation. Similarly, continued progress in handling image defects would be critical for processing the trillions of gigabytes envisioned by a bold proposal for ssEM imaging of an entire mouse brain (Advisory Committee to the NIH Director BRAIN Initiative Working Group 2. 2019).

## Methods

### Mouse

All procedures were in accordance with the Institutional Animal Care and Use Committees at the Baylor College of Medicine and the Allen Institute for Brain Science.

### Mouse line

Functional imaging was performed in a transgenic mouse expressing fluorescent GCaMP6f. For this data set, the mouse we used was a triple-heterozygote for the following three genes: (1) Cre driver: CamKIIa-Cre (Jax: 005359<https://www.jax.org/strain/005359>), (2) tTA driver: B6;CBA-Tg(Camk2a-tTA)1Mmay/J (Jax: 003010<https://www.jax.org/strain/003010>), (3) GCaMP6f Reporter: Ai93 (Allen Institute).

### Cranial window surgery

Anesthesia was induced with 3% isoflurane and maintained with 1.5% to 2% isoflurane during the surgical procedure. Mice were injected with 5-10 mg/kg ketoprofen subcutaneously at the start of the surgery. Anesthetized mice were placed in a stereotaxic head holder (Kopf Instruments) and their body temperature was maintained at 37°C throughout the surgery using a homeothermic blanket system (Harvard Instruments). After shaving the scalp, bupivicane (0.05 cc, 0.5%, Marcaine) was applied subcutaneously, and after 10-20 minutes an approximately 1 cm^2^ area of skin was removed above the skull and the underlying fascia was scraped and removed. The wound margins were sealed with a thin layer of surgical glue (VetBond, 3M), and a 13mm stainless-steel washer clamped in the headbar was attached with dental cement (Dentsply Grip Cement). At this point, the mouse was removed from the stereotax and the skull was held stationary on a small platform by means of the newly attached headbar. Using a surgical drill and HP 1/2 burr, a 3 mm craniotomy was made centered on the primary visual cortex (V1; 2.7mm lateral of the midline, contacting the lambda suture), and the exposed cortex was washed with ACSF (125mM NaCl, 5mM KCl, 10mM Glucose, 10mM HEPES, 2mM CaCl2, 2mM MgSO4). The cortical window was then sealed with a 3 mm coverslip (Warner Instruments), using cyanoacrylate glue (VetBond). The mouse was allowed to recover for 1-2 hours prior to the imaging session. After imaging, the washer was released from the headbar and the mouse was returned to the home cage.

### Widefield imaging

Prior to two-photon imaging, we acquired a low-magnification image of the 3mm craniotomy under standard illumination.

### Two-Photon imaging

Imaging for candidate mice was performed in V1, in a 400 ✕ 400 ✕ 200 µm volume with the superficial surface of the volume at the border of L1 and L2/3, approximately 100um below the pia. Laser excitation was at 920nm at 25-45mW depending on depth. The objective used was a 25✕ Nikon objective with a numerical aperture of 1.1, and the imaging point-spread function was measured with 500 nm beads and was approximately 0.5 ✕ 0.5 ✕ 3 µm^3^ in x, y, and z. Pixel dimensions of each imaging frame were 256 ✕ 256.

### Tissue preparation and staining

The protocol of Hua et al. (Hua, Laserstein, and Helmstaedter 2015) was combined with the protocol of Tapia et al. (Tapia et al. 2012) to accommodate a smaller tissue size and to improve TEM contrast. Mice were transcardially perfused with 2.5% paraformaldehyde and 1.25% glutaraldehyde. After dissection, 200 *μ*m thick coronal slices were cut with a vibratome and post-fixed for 12-48 h. Following several washes in CB (0.1 M cacodylate buffer pH 7.4), the slices were fixed with 2% osmium tetroxide in CB for 90 min, immersed in 2.5% potassium ferricyanide in CB for 90 min, washed with deionized (DI) water for 2 x 30 min, and treated with freshly made and filtered 1% aqueous thiocarbohydrazide at 40°C for 10 min. The slices were washed 2 ✕ 30 min with DI water and treated again with 2% osmium tetroxide in water for 30 min. Double washes in DI water for 30 min each were followed by immersion in 1% aqueous uranyl acetate overnight at 4°C. The next morning, the slices in the same solution were placed in a heat block to raise the temperature to 50°C for 2 h. The slices were washed twice in DI water for 30 min each, and then incubated in Walton’s lead aspartate pH 5.0 for 2 h at 50°C in the heat block. After another double wash in DI water for 30 min each, the slices were dehydrated in an ascending ethanol series (50%, 70%, 90%, 100% ✕ 3) 10 minutes each and two transition fluid steps of 100% acetonitrile for 20 min each. Infiltration with acetonitrile:resin dilutions (2p:1p, 1p:1p and 2p:1p) were performed on a gyratory shaker overnight for 4 days. Slices were placed in 100% resin for 24 hours followed by embedding in Hard Plus resin (EMS, Hatfield, PA). Slices were cured in a 60°C oven for 96 h. The best slice based on tissue quality and overlap with the 2p region was selected.

### Sectioning and collection

A Leica EM UC7 ultramicrotome and a Diatome 35 degree diamond ultra-knife were used for sectioning at a speed of 0.3 mm/sec. Eight to ten serial sections were cut at 40 nm thickness to form a ribbon, after which the microtome thickness setting was set to 0 in order to release the ribbon from the knife. Using an eyelash probe, pairs of ribbons were collected onto copper grids covered by 50 nm thick LUXEL film.

### Transmission electron microscopy

We made several custom modifications to a JEOL-1200EXII 120kV transmission electron microscope (Yin et al. 2019). A column extension and scintillator magnified the nominal field of view by tenfold with negligible loss of resolution. A high-resolution, large-format camera allowed fields-of-view as large as (13 µm)^2^ at 3.58 nm resolution. Magnification reduced the electron density at the phosphor, so a high-sensitivity sCMOS camera was selected and the scintillator composition tuned to generate high quality EM images with exposure times of 90-200 ms. Sections were acquired as a grid of 3840 x 3840 px images (“tiles”) with 15% overlap.

### Alignment in two blocks

The dataset was divided by sections into two blocks (1216 & 970 sections), with the first block containing substantially more folds. Initial alignment and reconstruction tests proceeded on the second block of the dataset. After achieving satisfactory results, the first block was added, and the whole dataset was further aligned to produce the final 3D image. The alignment process included stitching (assembling all tiles into a single image per section), rough alignment (aligning the set of section images with one affine per section), coarse alignment (nonlinear alignment on lower resolution data), and fine alignment (nonlinear alignment on higher resolution data).

### Alignment, block one

The tiles of the first block were stitched into one montaged image per section and rough aligned using a set of customized and automated modules based on the “TrakEM2” (Cardona et al. 2012) and “Render” (Zheng et al. 2018) software packages.

#### Stitching

After acquisition, a multiplicative intensity correction based on average pixel intensity was applied to the images followed by a lens distortion of individual tiles using non linear transformations (Kaynig et al. 2010). Once these corrections were applied, correspondences between tiles within a section were computed using SIFT features, and each tile was modeled with a rigid transform.

#### Rough alignment

Using 20x downsampled stitched images, neighboring sections were roughly aligned (Saalfeld et al. 2012). Correspondences were again computed using SIFT features, and each section was modeled with a regularized affine transform (90% affine + 10% rigid), and all correspondences and constraints were used to generate the final model of one affine transform per tile. These models were used to render the final stitched section image into rough alignment with block two.

### Alignment, block two

The second block was stitched and aligned using the methods of (Saalfeld et al. 2012) as implemented in Alembic (Macrina and Ih n.d.).

#### Stitching

For each section, tiles containing tissue without clear image defects were contrast normalized by centering the intensities at the same location in each tile, stretching the overall distribution between the 5th & 95th intensity percentiles. During imaging, a 20x downsampled overview image of the section was also acquired. Each tile was first placed according to stage coordinates, approximately translated based on normalized cross-correlation (NCC) with the overview image, and then finely translated based on NCC with neighboring tiles. Block matching was performed in the regions of overlap between tiles using NCC with 140 px block radius, 400 px search radius, and a spacing of 200 px. Matches were manually inspected with 1x coverage, setting per-tile-pair thresholds for peak of match correlogram, distance between first and second peaks of match correlograms, and correlogram covariance, and less frequently, targeted match removal. A graphical user interface was developed to allow the operator to fine-tune parameters on a section-by-section basis, so that a skilled operator completed inspection in 40 hrs. Each tile was modeled as a spring mesh, with nodes located at the center of each blockmatch operation, spring constants 1/100th of the constant for the between-tile springs, and the energy of all spring meshes within a section were minimized to a fractional tolerance of 10^-8^ using nonlinear conjugate gradient. The final render used a piecewise affine model defined by the mesh before and after relaxation, and maximum-intensity blending.

#### Rough alignment

Using 20x downsampled images, block matching between neighboring sections proceeded using NCC with 50 px block radius, 125 px search radius, and 250 px spacing. Matches were computed between nearest neighbor section pairs, then filtered manually in 8 hrs. Correspondences were used to develop a regularized affine model per section (90% affine + 10% rigid), which was rendered at full image resolution.

#### Coarse alignment

Using 4x downsampled images, NCC-based block matching proceeded 300 px block radius, 200 px search radius, and 500 px spacing. Matches were computed between nearest and next-nearest section pairs, then manually filtered by a skilled operator in 24 hrs. Each section was modeled as a spring mesh with spring constants 1/10th of the constant for the between-section springs, and the energy of all spring meshes within the block were minimized to a fractional tolerance of 10^-8^ using nonlinear conjugate gradient. The final render used a piecewise affine model defined by the mesh.

#### Fine alignment

Using 2x downsampled images, NCC-based block matching proceeded 200 px block radius, 113 px search radius, and 100 px spacing. Matches were computed between nearest and next-nearest section pairs, then manually filtered by a skilled operator in 24 hrs. Modeling and rendering proceeded as with coarse alignment, using spring constants were 1/20th of the constant for the between-section springs.

### Alignment, whole dataset

Blank sections were inserted manually between sections where the cutting thickness appeared larger than normal (11). The alignment of the whole dataset was further refined using the methods of (Saalfeld et al. 2012) as implemented in Alembic (Macrina and Ih n.d.).

#### Coarse alignment

Using 64x downsampled images, NCC-based block matching proceeded 128 px block radius, 512 px search radius, and 128 px spacing. Matches were computed between neighboring and next nearest neighboring sections, as well as 24 manually identified section pairs with greater separation, then manually inspected in 70 hrs. Section spring meshes had spring constants 1/20th of the constant for the between-section springs. Mesh relaxation was completed in blocks of 15 sections, 5 of which were overlapping with the previous block (2 sections fixed), each block relaxing to a fractional tolerance of 10^-8^. Rendering proceeded similarly as before.

#### Fine alignment

Using 4x downsampled images, NCC-based block matching proceeded 128 px block radius, 512 px search radius, and 128 px spacing. Matches were computed between the same section pairs as in coarse alignment. Matches were excluded only be heuristics. Modeling and rendering proceeded similar to coarse alignment, with spring constants 1/100th the constant for the between-section springs. Rendered image intensities were linearly rescaled in each section based on the 5th and 95th percentile pixel values.

### Image volume estimation

The imaged tissue has a trapezoidal shape in the sectioning plane. Landmark points were placed in the aligned images to measure this shape. We report cuboid dimensions for simplicity and comparison using the trapezoid midsegment length. The original trapezoid has a short base length of 216.9μm, long base length of 286.2μm, and height 138.3μm. The imaged data has 2176 sections, which measures 87.04μm with a 40nm slice thickness.

### Image defect handling

Cracks, folds, and contaminants were manually annotated as binary masks on 256x downsampled images, dilated by 2 px, then inverted to form a defect mask. A tissue mask was created using nonzero pixels in the 256x downsampled image, then eroded by 2 px to exclude misalignments at the edge of the image. The image mask is the union of the tissue and defect masks, and it was upsampled and applied during the final render to set pixels not included in the mask to zero. We created a segmentation mask by excluding voxels that had been excluded by the image mask for three consecutive sections. The segmentation mask was applied after affinity prediction to set affinities not included in the mask to zero.

### Affinity prediction

Human experts used [VAST](https://software.rc.fas.harvard.edu/lichtman/vast/) (Berger, Seung, and Lichtman 2018) to manually segment multiple subvolumes from the current dataset and a similar dataset from mouse V1. Annotated voxels totaled 1.29 billion at full image resolution.

We trained a 3D convolutional network to generate three nearest neighbor (Turaga et al. 2010) and 13 long-range affinity maps (K. Lee et al. 2017). Each long-range affinity map was constructed by comparing an equivalence relation (Jain, Sebastian Seung, and Turaga 2010) of pairs of voxels spanned by an “offset” edge (to preceding voxels at distances of 4, 8, 12, and 16 in x and y, and 2, 3, 4 in z). Only the nearest neighbor affinities were used beyond inference time; long-range affinities were used solely for training. The network architecture was modified from the “Residual Symmetric U-Net” of Lee et al. (K. Lee et al. 2017). We trained on input patches of size 128×128×20 at 7.16×7.16×40 nm^3^ resolution. The prediction during training was bilinearly upsampled to full image resolution before calculating the loss.

Training utilized synchronous gradient updates computed by four Nvidia Titan X Pascal GPUs each with a different input patch. We used the AMSGrad variant (Reddi, Kale, and Kumar 2019) of the Adam optimizer (Kingma and Ba 2014), with PyTorch’s default settings except step size parameter α = 0.001. We used the binary cross-entropy loss with “inverse margin” of 0.1 (Huang and Jain 2013); patch-wise class rebalancing (K. Lee et al. 2017) to compensate for the lower frequency of boundary voxels; training data augmentation including flip/rotate by 90°, brightness and contrast perturbations, warping distortions, misalignment/missing section simulation, and out-of-focus simulation (K. Lee et al. 2017); and lastly several new types of data augmentation including the simulation of lost section and co-occurrence of misalignment/missing/lost section. Distributed computation of affinity maps used chunkflow (Wu et al. 2019). The computation was done with images at 7.16×7.16×40 nm^3^ resolution. The whole volume was divided into 1280×1280×140 chunks overlapping by 128×128×10, and each chunk was processed as a task. The tasks were injected into a queue (Amazon Web Service Simple Queue Service). For 2.5 days, 1000 workers (Google Cloud n1-highmem-4 with 4 vCPUs and 26 GB RAM, deployed in Docker image using Kubernetes) fetched and executed tasks from the queue as follows. The worker read the corresponding chunk from Google Cloud Storage using CloudVolume (seung-lab n.d.), and applied previously computed masks to black out regions with image defects. The chunk was divided into 256×256×20 patches with 50% overlap. Each patch was processed to yield an affinity map using PZNet, a CPU inference framework (Popovych et al. 2020). The overlapping output patches were multiplied by a bump function, which weights the voxels according to the distance from patch center, for smooth blending and then summed. The result was cropped to 1024×1024×120 vx and then previously computed segmentation masks were applied (see Image defect handling above).

### Watershed and size-dependent single linkage clustering

The affinity map was divided into 514×514×130 chunks that overlapped by 2 voxels in each direction. For each chunk we ran a watershed and clustering algorithm (Zlateski and Seung 2015) with special handling of chunk boundaries. If the descending flow of watershed terminated prematurely at a chunk boundary, the voxels around the boundary were saved to disk so that domain construction could be completed later on. Decisions about merging boundary domains were delayed, and information was written to disk so decisions could be made later. After the chunks were individually processed, they were stitched together in a hierarchical fashion. Each level of the hierarchy processed the previously delayed domain construction and clustering decisions in chunk interiors. Upon reaching the top of the hierarchy, the chunk encompassed the entire volume, and all previously delayed decisions were completed.

### Mean affinity agglomeration

The watershed supervoxels and affinity map were divided into 513×513×129 chunks that overlapped by 1 in each direction. Each chunk was processed using mean affinity agglomeration (K. Lee et al. 2017; Funke et al. 2019). Agglomeration decisions at chunk boundaries were delayed, and information about the decisions was saved to disk. After the chunks were individually processed, they were combined in a hierarchical fashion similar to the watershed process.

### Synaptic cleft detection

Synaptic clefts were annotated by human annotators within a 310.7μm^3^ volume, which was split into 203.2μm^3^ training, 53.7μm^3^ validation, and 53.7μm^3^ test sets. We trained a version of the Residual Symmetric U-Net (K. Lee et al. 2017) with 3 downsampling levels instead of 4, 90 feature maps at the 3rd downsampling instead of 64, and ‘resize’ upsampling rather than strided transposed convolution. Images and labels were downsampled to 7.16×7.16×40nm^3^ image resolution. To augment the training data, input patches were transformed by (1) introducing misalignments of up to 17 pixels, (2) blacking out up to 5 sections, (3) blurring up to 5 sections, (4) warping, (5) varying brightness and contrast, and (6) flipping and rotating by multiples of 90 degrees. Training used PyTorch (Adam et al. 2017), and the Adam optimizer (Kingma and Ba 2014). The learning rate started from 10^−3^, and was manually annealed three times (505k training updates), before adding 67.2μm^3^ of extra training data for another 670k updates. The extra training data focused on false positive examples from the network’s predictions at 505k training updates, mostly around blood vessels. The trained network achieved 93.0% precision and 90.9% recall in detecting clefts of the test set. This network was applied to the entire dataset using the same distributed inference setup as affinity map inference. Connected components of the thresholded network output that were at least 50 voxels at 7.16×7.16×40nm^3^ resolution were retained as predicted synaptic clefts.

### Synaptic partner assignment

Presynaptic and postsynaptic partners were annotated for 387 clefts, which were split into 196, 100, and 91 examples for training, validation, and test sets. A network was trained to perform synaptic partner assignment via a voxel association task(Turner, Lee, et al. 2020). Architecture and augmentations were the same as for the synaptic cleft detector. Test set accuracy was 98.9% after 710k training iterations. The volume was separated into non-overlapping chunks of size 7.33μm x 7.33μm x 42.7μm (1024 x 1024 x 1068 voxels), and the net was applied to each cleft in each chunk. This yielded a single prediction for interior clefts. For a cleft that crossed at least one boundary, we chose the prediction from the chunk which contained the most voxels of that cleft. Cleft predictions were merged if they connected the same synaptic partners and their centers-of-mass were within 1 μm. This resulted in 3,556,643 final cleft predictions.

### PyC proofreading

The mean affinity graph of watershed supervoxels was stored in our PyChunkedGraph backend, which uses an octree to provide spatial embedding for fast updates of the connected component sets from local edits. We modified the Neuroglancer frontend (Maitin-Shepard 2019) to interface with this backend so users directly edit the agglomerations by adding and removing edges in the supervoxel graph (merge and split agglomerations). Connected components of this graph are meshed in chunks of supervoxels, and chunks affected by edits are updated in real-time so users can always see a 3D representation of the current segmentation. Using a keypoint for each object (e.g. soma centroid), objects are assigned the unique ID of the connected component for the supervoxel which contains that location. This provides a means to update the object’s ID as edits are made.

Cell bodies in the EM volume were semi-automatically identified. Pyramidal cells were identified by morphological features, including density of dendritic spines, presence of apical and basal dendrites, direction of main axon trunk, and cell body shape. We selected a subset of the 417 PyCs for proofreading based on the amount of visible neurite within the volume. A team of annotators used the meshes to detect errors in dendritic trunks and axonal arbors, then to correct those errors with 50,000 manual edits in 1,044 person-hours. After these edits, pyramidal cells were skeletonized, and both the branch and end points of these skeletons were identified automatically (with false negative rates of 1.7% and 1.4%, as estimated by annotators). Human annotators reviewed each point to ensure no merge errors and extend split errors where possible (210 person-hours). Putative broken spines targeted by PyCs were identified by selecting objects that received one or two synapses. Annotators reviewed, and attached these with 174 edits in 24 person-hours. Some difficult mergers came from small axonal boutons merged to dendrites. We identified these cases by inspecting any predicted presynaptic site that resided within 7.5 μm of a postsynaptic site of the same cell, and corrected them with 50 person-hours.

### Estimation of final error rates

After proofreading was complete, a single annotator inspected 12 pyramidal cells and spent 18 hrs to identify all remaining errors in dendritic trunks and axonal arbors. The PyC proofreading protocol was designed to correct all merge errors, though not necessarily correct split errors caused by a masked segmentation. So this error estimation includes all merge errors identified and only split errors caused by less than 3 consecutive sections of masked segmentation. For 18.7 mm of dendritic path length inspected, 3 false splits (falsely excluding 160 synapses) and 3 false merges (falsely including 117 synapses) were identified (99% precision and 99% recall for incoming synapses). For 3.6 mm of axonal path length inspected, 2 false splits (falsely excluding 4 synapses) and 1 false merge (falsely including 9 synapses) were identified (98% precision and 99% recall for outgoing synapses). We also sampled four dendritic branches with a collective 0.7 mm of path length, and identified 126 false negative and 0 false positive spines (88% recall of spines).

### PyC-PyC synapse proofreading

Synapses between PyC were extracted from the automatically detected and assigned synapses. We reviewed these synapses manually with 2x redundancy (1972 correct synapses out of 2433 putative synapses). 2 predicted synapses out of these were “merged” with other synaptic clefts. These cases were excluded from further analysis. 1 synapse was incorrectly assigned to a PyC and removed from the analysis. 1 other synapse was “split” into two predictions, and these predictions were merged for analysis. We were not able to calculate spine head volumes for 8 out of these 1968 synapses and they were excluded from the analysis. This left 1960 synapses admitted into the analysis.

### Synapses from other excitatory axons

We randomly sampled synapses onto the PyCs and evaluated whether they are excitatory or inhibitory based on their shape, appearance and targeted compartment (n=881 single excitatory synapses). We randomly sampled connections of two synapses onto PyCs and evaluated whether their presynaptic axon is excitatory or inhibitory and checked for reconstruction errors. Here, we manually checked that the automatically reconstructed path between the two synapses along the 3D mesh of the axon was error free (n=446 pairs of excitatory synapses). Those axons were allowed to contain errors elsewhere and we did not proofread any axons to obtain these pairs.

### Dendritic spine heads

We extracted a 7.33 x 7.33 x 4 μm^3^ cutout around the centroid of each synapse. The postsynaptic segment within that cutout was skeletonized using kimimaro (https://github.com/seung-lab/kimimaro), yielding a set of paths traveling from a root node to each leaf. The root node was defined as the node furthest from the synapse coordinate. Skeleton nodes participating in fewer than three paths were labeled as “spine” while others were labeled as “shaft.” The shaft labels were dilated along the skeleton until either (1) the distance to the segment boundary of the next node was more than 50 nm less than that of the closest (shaft) branch point, or (2) dilation went 200 nodes beyond the branch point. Each synapse was associated with its closest skeleton node, and a contiguous set of “spine” labeled nodes. We finally separated spine head from neck by analyzing the distance to the segment boundary (DB) moving from the root of the spine to the tip. After segmenting the spine from the rest of the segment, we chose two anchor points: (1) the point with minimum DB value across the half of the spine towards the dendritic shaft, and (2) the point with maximum DB value across the other half. A cut point was defined as the first skeleton node moving from anchor 1 to anchor 2 whose DB value was greater than ⅓ DB_anchor1_ + ⅔ DB_anchor2_. Accounting for slight fluctuations in the DB value, we started the scan for the cut point at the closest node to anchor 2 that had DB value less than ⅕ DB_anchor1_ + ⅘ DB_anchor2_. The skeleton of the spine head was defined as the nodes beyond this cut point to a leaf node, and the spine head mesh was defined as all spine mesh vertices which were closest to the spine head skeleton. The mesh of each head was identified as the subset of the postsynaptic segment mesh whose closest skeleton node was contained within the nodes labeled as spine head. We then estimated the volume of this spine head by computationally sealing this mesh and computing its volume.

We identified poor extractions by computing the distance between each synapse centroid and the nearest node of its inferred spine head mesh. We inspected each inferred spine head for which this distance was greater than 35 nm, and corrected the mesh estimates of mistakes by relabeling mesh vertices using a 3D voronoi tessellation of points placed by a human annotator. **Endoplasmic reticulum.** We manually evaluated all spine heads between PyCs admitted to the analysis for whether they contained a spine apparatus (SA), endoplasmic reticulum that is not an SA (ER) or none (no ER). We required the presence of at least two (usually parallel) membrane saccules for SA. Dense plate/region (synaptopodin and actin) in between membrane saccules was an indicator. We found SA in spine heads and spine necks. We considered single lumens of organelles connecting to the ER network in the shaft as ER. We required that every ER could be traced back to the ER network in the dendritic shaft.

### Mixture and hidden Markov models

Spine volumes and synapse sizes were log_10_-transformed before statistical modeling. Maximum likelihood estimation for a binary mixture of normal distributions used the expectation-maximization algorithm as implemented by Pomegranate (Schreiber 2017). The algorithm was initialized using the k-means algorithm with k=2. For cleft size, the normal distributions were truncated at a lower bound of log_10_(50) voxels, the cutoff used in cleft detection. The truncation was implemented by modifications to the source code of Pomegranate. (A similar truncation was used when fitting a single normal distribution to cleft sizes.) The joint distribution at dual connections was fit by hidden Markov Models (HMMs) with two latent states and emission probabilities given by normal or truncated normal distributions. Hidden Markov models are trained on ordered pairs, yet each dataset used for training was reflected to contain both orders of each synapse pair.

### Parametric test for bimodality

For binary mixtures of normal distributions, the parameter regimes for bimodal and unimodal behaviors are known (Robertson and Fryer 1969). The likelihood ratio of the best-fitting bimodal and unimodal models can be used for model selection (Holzmann and Vollmer 2008). Mixture models were fit using Sequential Least Squares Programming using constraints on the parameter regimes for unimodal fits. We computed *p*-values using Chernoff’s extension to boundary points of hypothesis sets (Chernoff 1954) of Wilks’ theorem governing asymptotics of the likelihood ratio (Wilks 1938).

### Correlation analysis

We assigned state probabilities to each dual synaptic pair using the best fit HMM. The following was done for SS and LL states independently. In each sampling iteration (n=10,000) we assigned individual synapse pairs to the state in question based on independent biased coin flips weighted by their respective state probability. For every such obtained sample we computed the pearson correlation. For visualization in Fig. 4g,h we applied a kernel density estimation (bw=0.15 in log10-space).

### Skeletonization

We developed a skeletonization algorithm similar to (Sato et al. 2000) that operates on meshes. For each connected component of the mesh graph, we identify a root and find the shortest path to the farthest node. This procedure is repeated after invalidating all mesh nodes within the proximity of the visited nodes until no nodes are left to visit. We make our implementation available through our package MeshParty (https://github.com/sdorkenw/MeshParty).

### Estimation of path lengths

We skeletonized all PyCs and labeled their first branch points close to the soma according to the compartment type of the downstream branches (axon, dendrite, ambiguous). If no branch point existed in close proximity a point at similar distance was placed. All skeleton nodes downstream from these nodes seen from the soma were labeled according to these labels. This allowed us to estimate path lengths for each compartment with the path up to the first branch point labeled as perisomatic (axon: 100 mm, dendrite: 520 mm, perisomatic: 40 mm, ambiguous: 10 mm). We estimated that our skeletons were overestimated by 11% due to following the mesh edges and corrected all reported pathlengths accordingly.

## Data availability

The raw images, segmentation, and synaptic connectivity will be made available upon or before publication.

## Code availability

All software is open source and available at http://github.com/seung-lab if not otherwise mentioned.

Alembic: Stitching and alignment.

CloudVolume: Reading and writing volumetric data, meshes, and skeletons to and from the cloud

Chunkflow: Running convolutional nets on large datasets

DeepEM: Training convolutional nets to detect neuronal boundaries. DynamicAnnotationFramework: Proofreading and connectome updates (visit https://github.com/seung-lab/AnnotationPipelineOverview for repository list) Igneous: Coordinating downsampling, meshing, and data management.

MeshParty: Interaction with meshes and mesh-based skeletonization (https://github.com/sdorkenw/MeshParty)

MMAAPP: Watershed, size-dependent single linkage clustering, and mean affinity agglomeration.

PyTorchUtils: Training convolutional nets for synapse detection and partner assignment (https://github.com/nicholasturner1/PyTorchUtils).

Synaptor: Processing output of the convolutional net for predicting synaptic clefts (https://github.com/nicholasturner1/Synaptor).

TinyBrain and zmesh: Downsampling and meshing (precursors of the libraries that were used).

## Supplementary information

Supplementary information is available for this paper.

## Acknowledgements

Supported by the Intelligence Advanced Research Projects Activity (IARPA) via Department of Interior/ Interior Business Center (DoI/IBC) contract numbers D16PC00003, D16PC00004, and D16PC0005. The U.S. Government is authorized to reproduce and distribute reprints for Governmental purposes notwithstanding any copyright annotation thereon. HSS also acknowledges support from NIH/NINDS U19 NS104648, ARO W911NF-12-1-0594, NIH/NEI R01 EY027036, NIH/NIMH U01 MH114824, NIH/NINDS R01NS104926, NIH/NIMH RF1MH117815, and the Mathers Foundation, as well as assistance from Google, Amazon, and Intel. We thank S. Koolman, M. Moore, S. Morejohn, B. Silverman, K. Willie, and R. Willie for their image analyses, Garrett McGrath for computer system administration, and May Husseini and Larry and Janet Jackel for project administration. We are grateful to J. Maitin-Shepard for neuroglancer and P. H. Li and V. Jain for helpful discussions. We thank D. W. Tank, K. Li, Y. Loewenstein, J. Kornfeld, A. Wanner, M. Tsodyks, D. Markowitz and G. Ocker for advice and feedback. We thank the Allen Institute for Brain Science founder, Paul G. Allen, for his vision, encouragement and support. Disclaimer: The views and conclusions contained herein are those of the authors and should not be interpreted as necessarily representing the official policies or endorsements, either expressed or implied, of IARPA, DoI/IBC, or the U.S. Government.

## Contributions

EF, JR, and AST performed surgery on the mouse and calcium-imaging of neural activity. JB, MT, and NMC prepared the sample for electron microscopy. ALB, AAB, DJB, JB, and NMC generated the electron microscopy dataset. RT, GM, and YL stitched and rough aligned 1216 sections. DI and TM stitched and rough, coarse, & fine aligned 970 sections. TM coarse & fine aligned all sections. WW built the task manager for distributed alignment. WMS wrote the software for reading and writing data in cloud storage, initially in collaboration with IT. TM supervised ground truth annotations. KL trained the convolutional network for boundary detection with help from JZ. SP implemented a framework for CPU-based 3D convolutional net inference with help from AZ. JW used the convolutional net to generate the affinity map. RL segmented the affinity map with help from AZ. NLT trained the convolutional nets for synapse detection and partner assignment. JW used the convolutional net to produce a synapse map. NLT segmented the synapse map and applied the partner assignment net to the dataset. SD, NK, and JZ created the proofreading system. SCY, TM, SD, and AMW supervised proofreading and annotation efforts. FC, ALB, NMC, SCY, SD and CSM contributed proofreading and annotations. NK and MC adapted and improved Neuroglancer for proofreading and annotations and, together with WMS, meshed the segmentation. CJ built the early data sharing system. SD, FC, and CSM designed and developed the connectome versioning system. SD, NLT, and TM analyzed the synaptic connectivity with input from HSS, FC, NMC, CSM, and RCR. HSS and SD wrote the paper, with contributions from NLT, TM, NK, KL, MT, NCM and help from FC, CSM. SS and LB managed the project at the Allen Institute. TM managed the reconstruction team. HSS, RCR, NMC, JR, and AST managed the multi-institution collaboration.

## Author information

These authors contributed equally: Sven Dorkenwald, Nicholas L. Turner, Thomas Macrina, Kisuk Lee, Ran Lu, Jingpeng Wu, Agnes L. Bodor, Adam A. Bleckert, Derrick Brittain

---

**Extended Data Figure 1.**
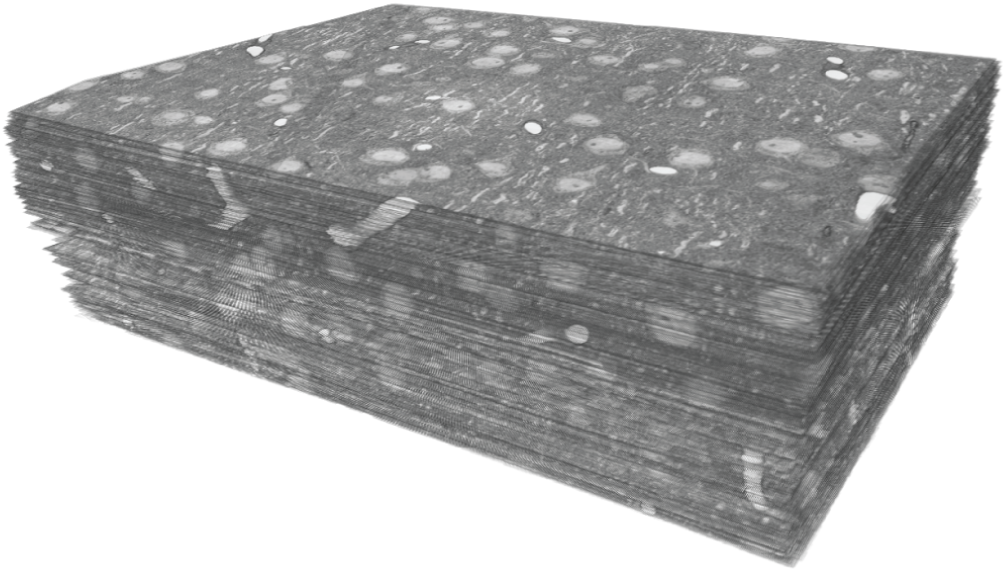
Reconstruction of connections between L2/3 pyramidal cells. 250×140×90 µm^3^ 3D image stack from layer 2/3 of mouse primary visual cortex

**Extended Data Figure 2.**
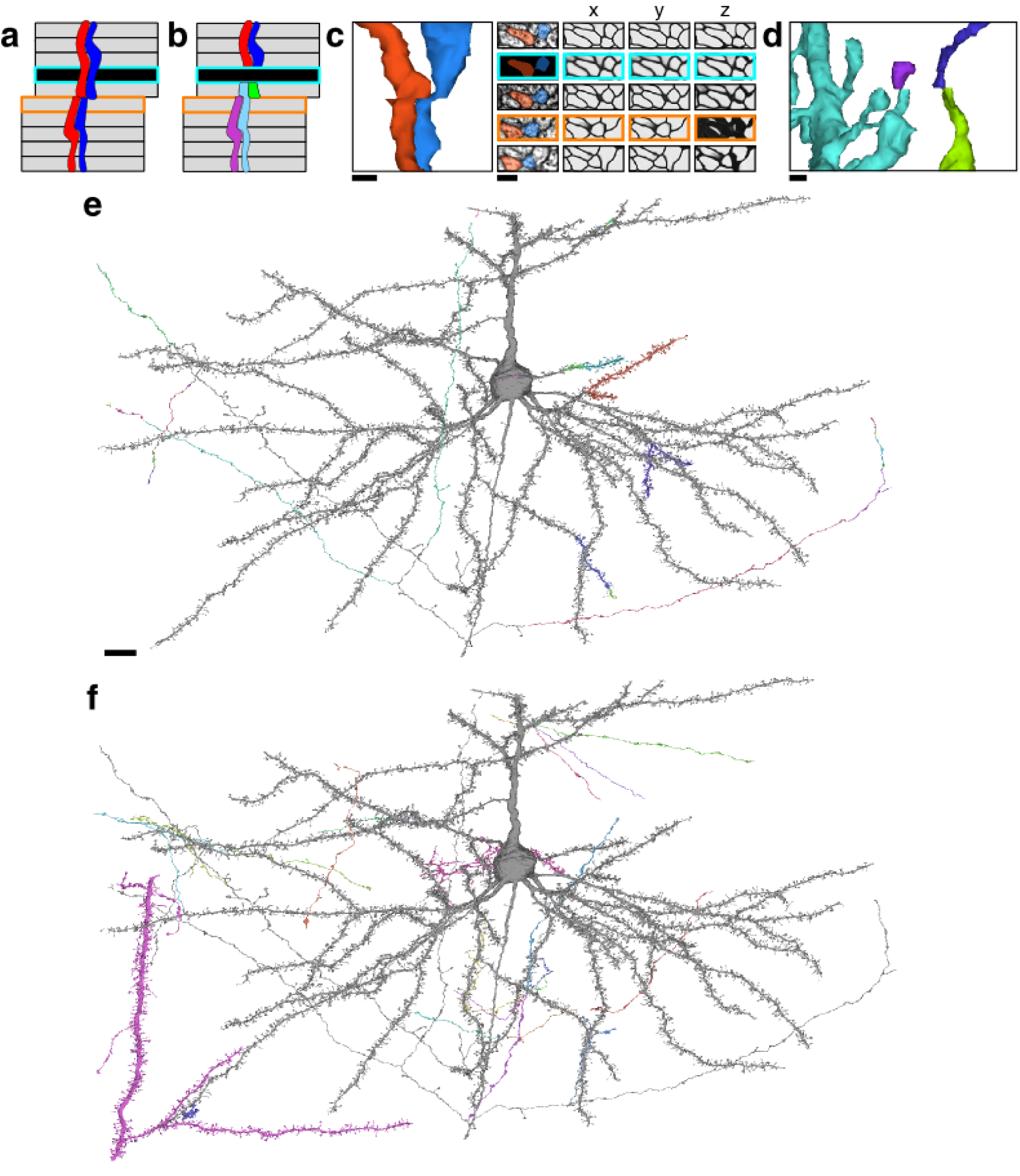
Examples of reconstructed neurites near image defects. **(a)** Illustration of a possible pair of neurites that pass through both missing section (cyan) and misalignment (orange) defects. **(b)** Illustration of a naive segmentation of the pair in (a). **(c)** Same examples as in Fig. 1, accompanied by affinity map. Scale bar 300 nm. **(d)** Near a larger misalignment, the displacement is larger than the width of a thin neurite, and the convolutional net is unable to trace through the misalignment. Scale bar 300nm. **(e)** A proofread neuron (gray) with segments merged during proofreading (colored). Scale bar 10 μm. **(f)** The same proofread neuron in (e) with pieces split during proofreading (colored).

**Extended Data Figure 3.**
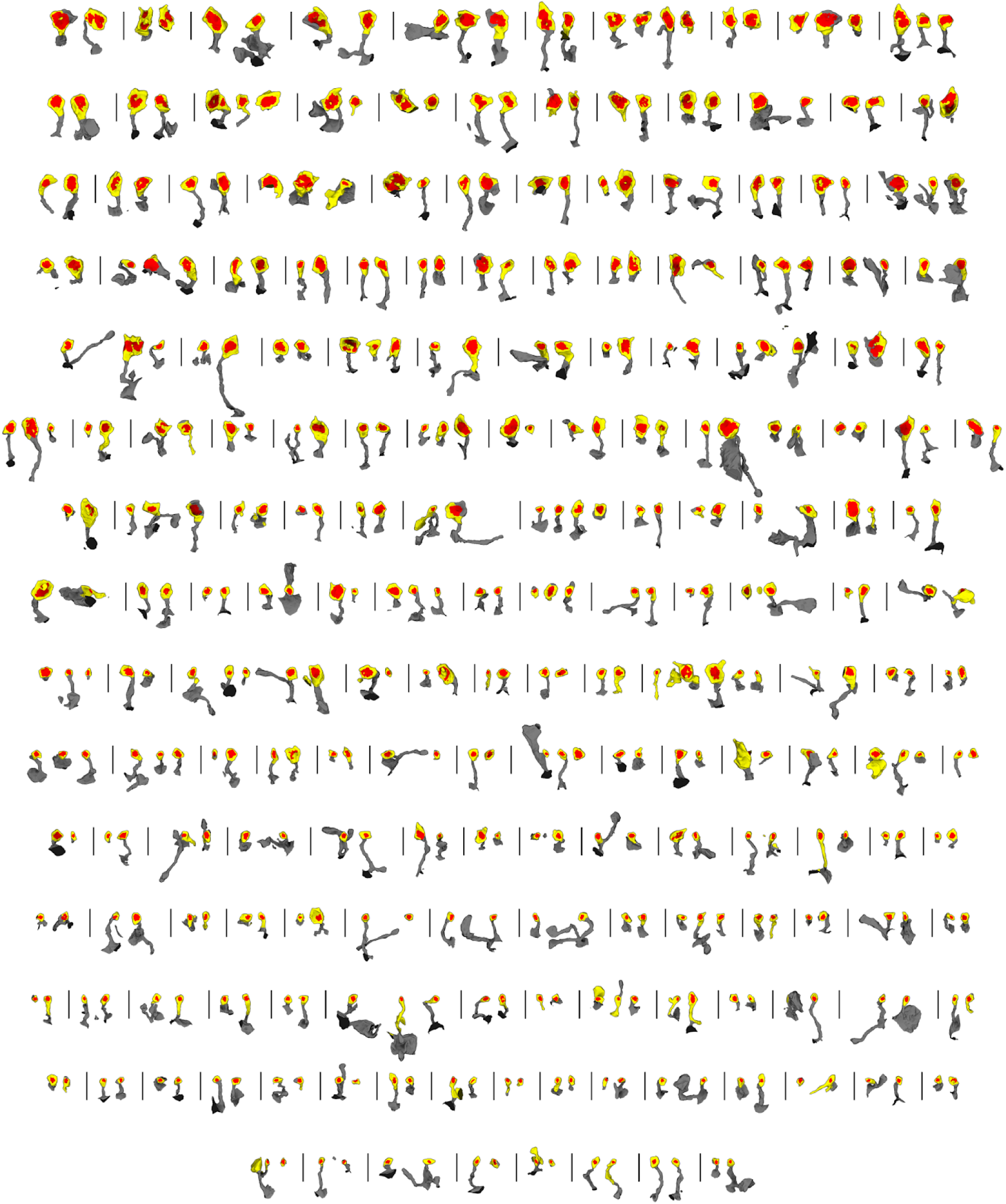
Renderings of all synapses from multisynaptic connections between L2/3 pyramidal cells. Dendritic spines (yellow) and synaptic clefts (red) are rendered in 3D. Most are dual connections (160), but there are also triples (24), quadruples (3), and quintuples (2).

**Extended Data Figure 4.**
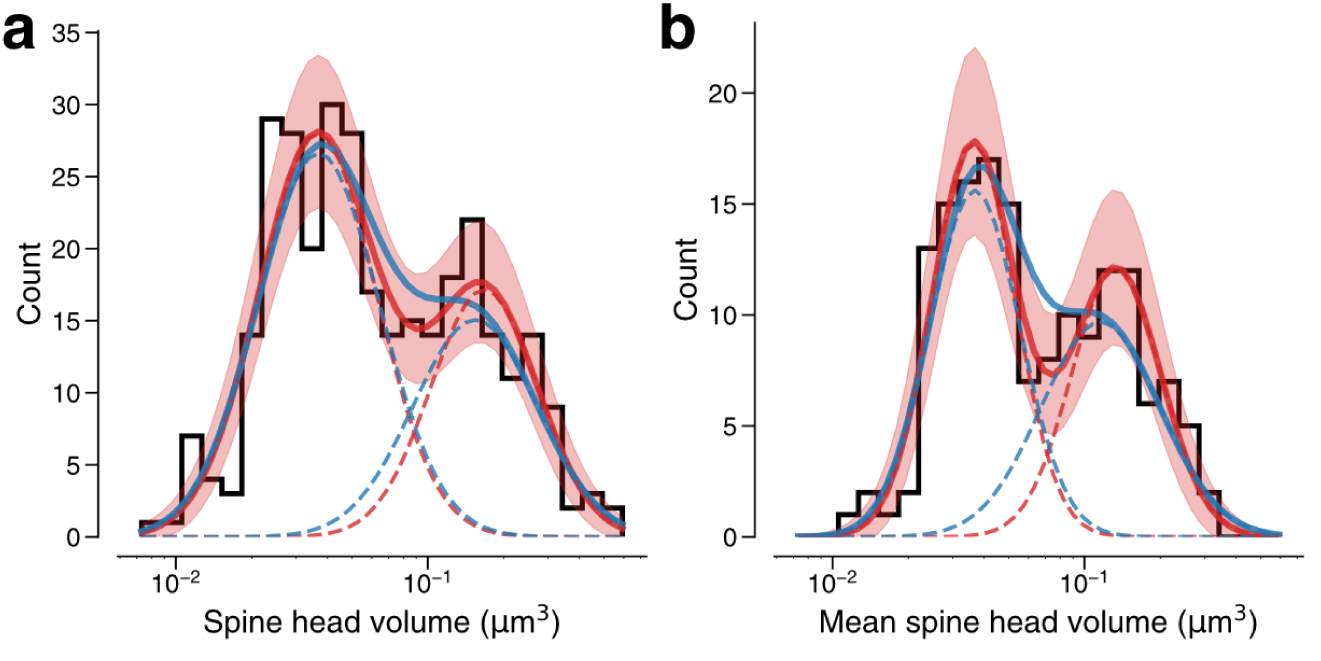
Modeling spine volume with a bimodal versus unimodal mixture of two normal distributions. **(a)** Spine volumes belonging to dual connections between L2/3 pyramidal cells. A bimodal mixture (red, solid) of two normal distributions (red, dashed) is a better fit than a unimodal mixture (blue, solid) of two normal distributions (blue, dashed) (likelihood ratio test, p=0.0425, n=320). The bimodal mixture weights are 60:40. **(b)** Dual connections between L2/3 pyramidal cells, each summarized by the geometric mean of spine volumes. A bimodal mixture (red, solid) of two normal distributions (red, dashed) is a better fit than a unimodal mixture (blue, solid) of two normal distributions (blue, dashed) (likelihood ratio test, p=0.0059, n=160). Error bars are ± 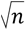 of the model fit.

**Extended Data Figure 5.**
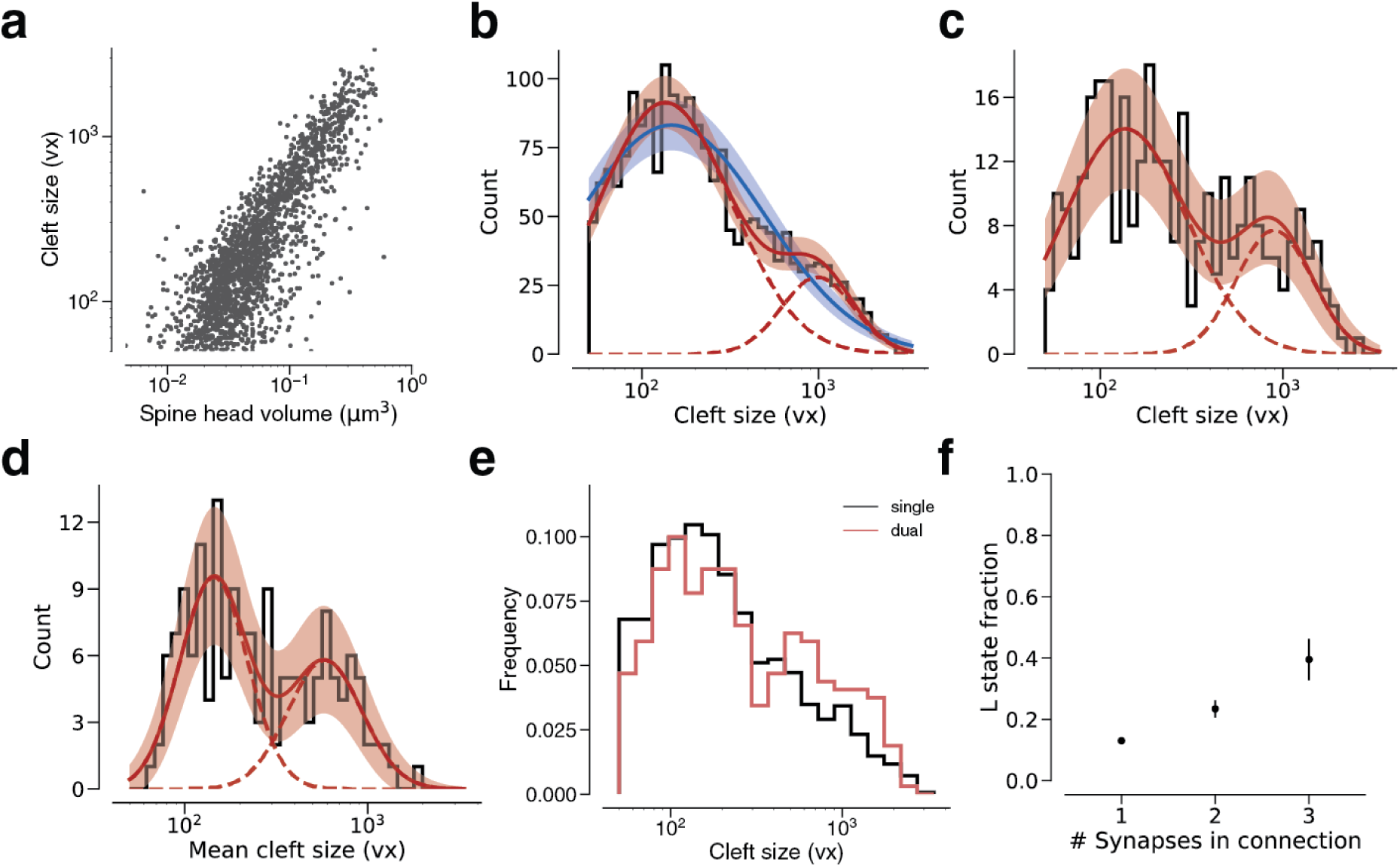
Modeling cleft size with a mixture of two normal distributions. **(a)** Spine volume vs. cleft size for all L2/3 PyC - L2/3 PyC synapses. **(b)** Histogram of same spine volumes, logarithmic scale. A mixture (red, solid) of two normal distributions (red, dashed) fits better (likelihood ratio test, p<1e-63, n=1960) than a single normal (blue). **(c)** Cleft sizes belonging to dual connections between L2/3 PyCs, modeled by a mixture (red, solid) of two normal distributions (red, dashed, likelihood ratio test, p=0.02, n=320). **(d)** Dual connections between L2/3 PyCs, each summarized by the geometric mean of two cleft sizes, modeled by a mixture (red, solid) of two normal distributions (red, dashed, likelihood ratio test, p=0.037, n=160). **(e)** Comparison of cleft sizes for single (black) and dual (red) connections. **(f)** Probability of the “L” state (mixture weight) versus number of synapses in the connection. Error bars are ± 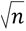 of the model fit.

**Extended Data Figure 6:**
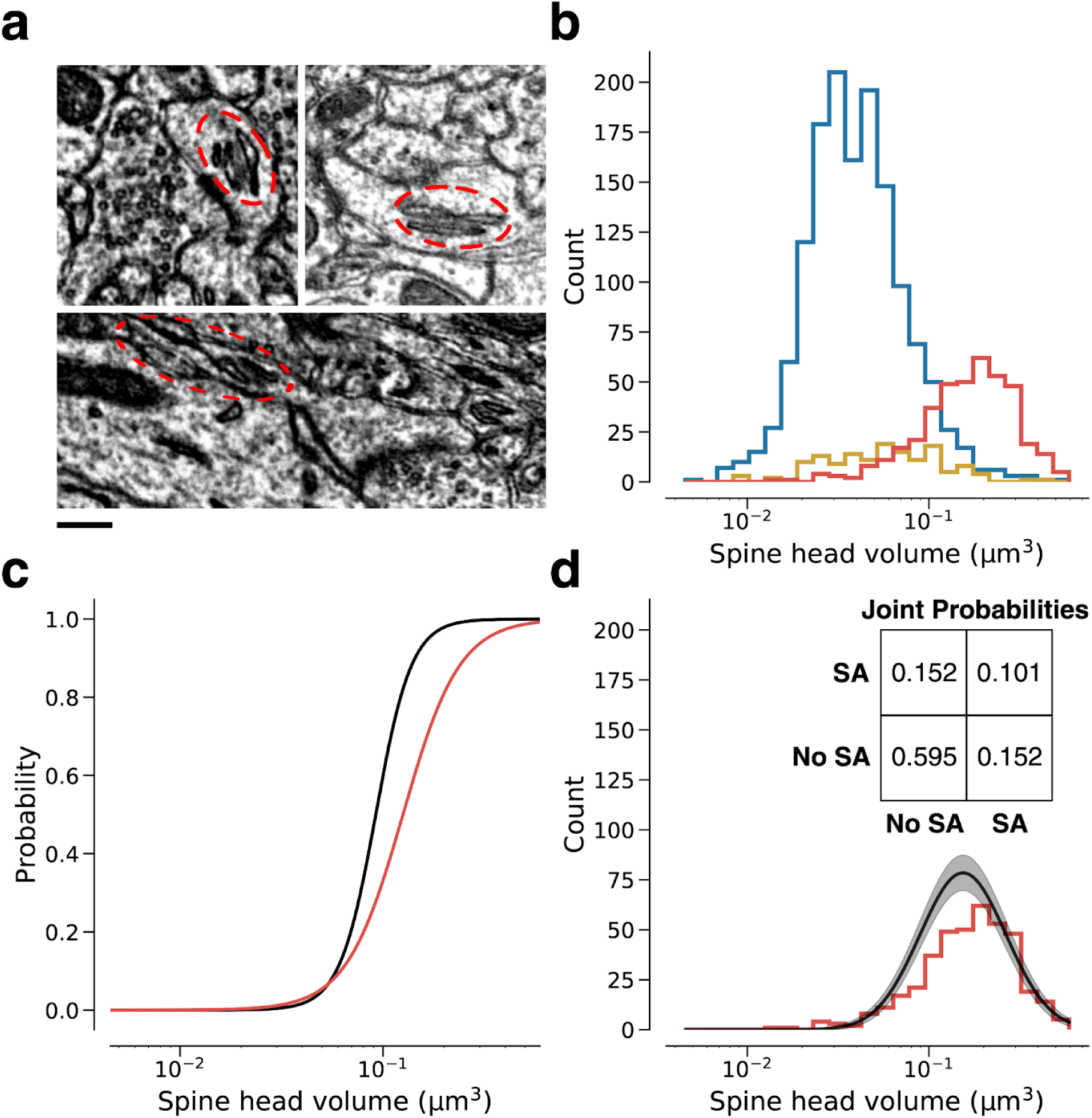
Relation of dendritic spine volume to spine apparatus. **(a)** Examples of spine apparatus (SA) in EM images. Scale bar 300nm. **(b)** Spine volume distributions of synapses with no ER (blue), smooth ER (yellow), and SA (red). **(c)** Likelihood of SA (red) and “L” state (black) conditioned on spine volume. **(d)** Spine volume distribution conditioned on SA (red) compared with size distribution conditioned on “L” state (black). Inset: Joint probability distribution of SA within dual connections. Error bars are ± 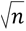 of the model fit.

**Extended Data Figure 7.**
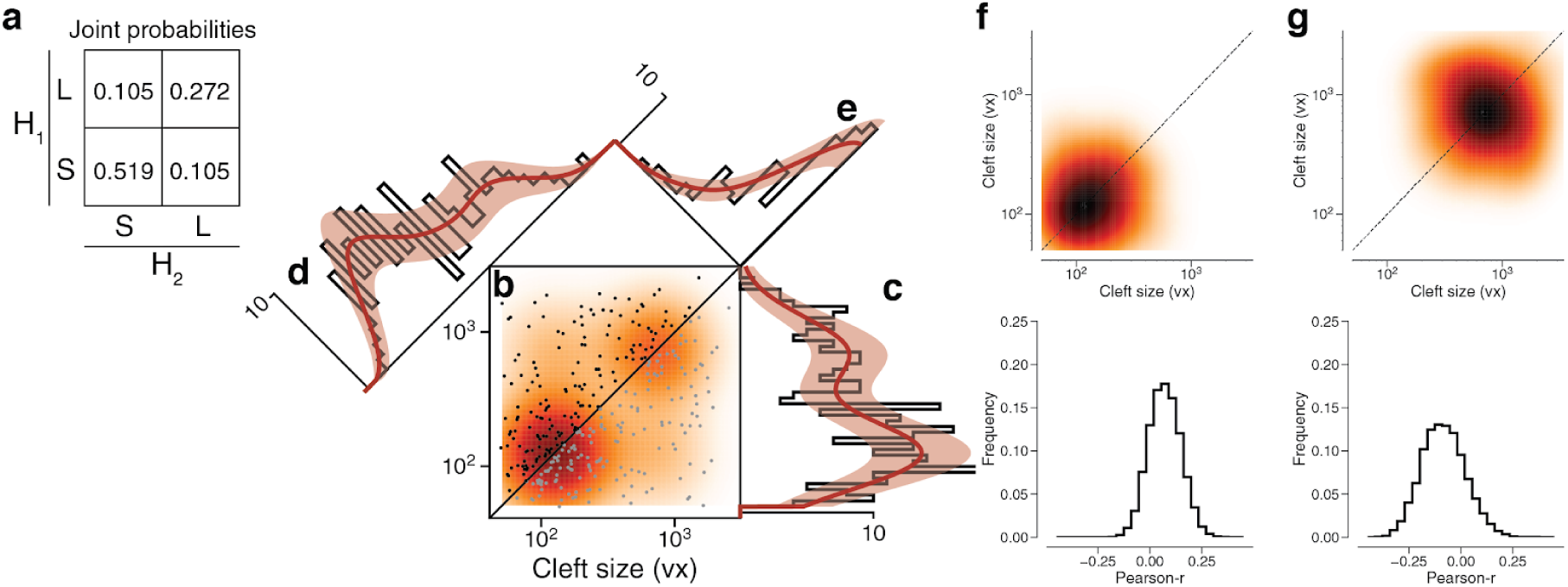
Latent state correlations between clefts at dual connections. **(a)** The latent states of synapses in a dual connection are positively correlated with each other. The latent states are more likely to be the same (SS or LL) rather than different (SL or LS), as shown by the joint probability distribution. **(b)** Scatter plot of cleft sizes (black, lexicographic ordering) for dual connections between L2/3 pyramidal cells. Scatter plot points are mirrored across the diagonal (gray). The joint distribution is fit by a mixture model (orange) like that of Fig. 3f, but with latent states that are correlated as described below. **(c)** Projecting the points onto the vertical axis yields a histogram of cleft sizes for dual connections, the same as in Fig. 3d. Model is derived from the joint distribution. **(d)** Projecting the points onto the x=y diagonal yields a histogram of the geometric mean of cleft sizes for dual connections, the same as in Fig. 3e. Model is derived from the joint distribution. **(e)** Projecting the points onto the x=−y diagonal yields a histogram of the ratio of cleft sizes for dual connections. Error bars are ± 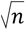 of the model fit.

**Extended Data Figure 8.**
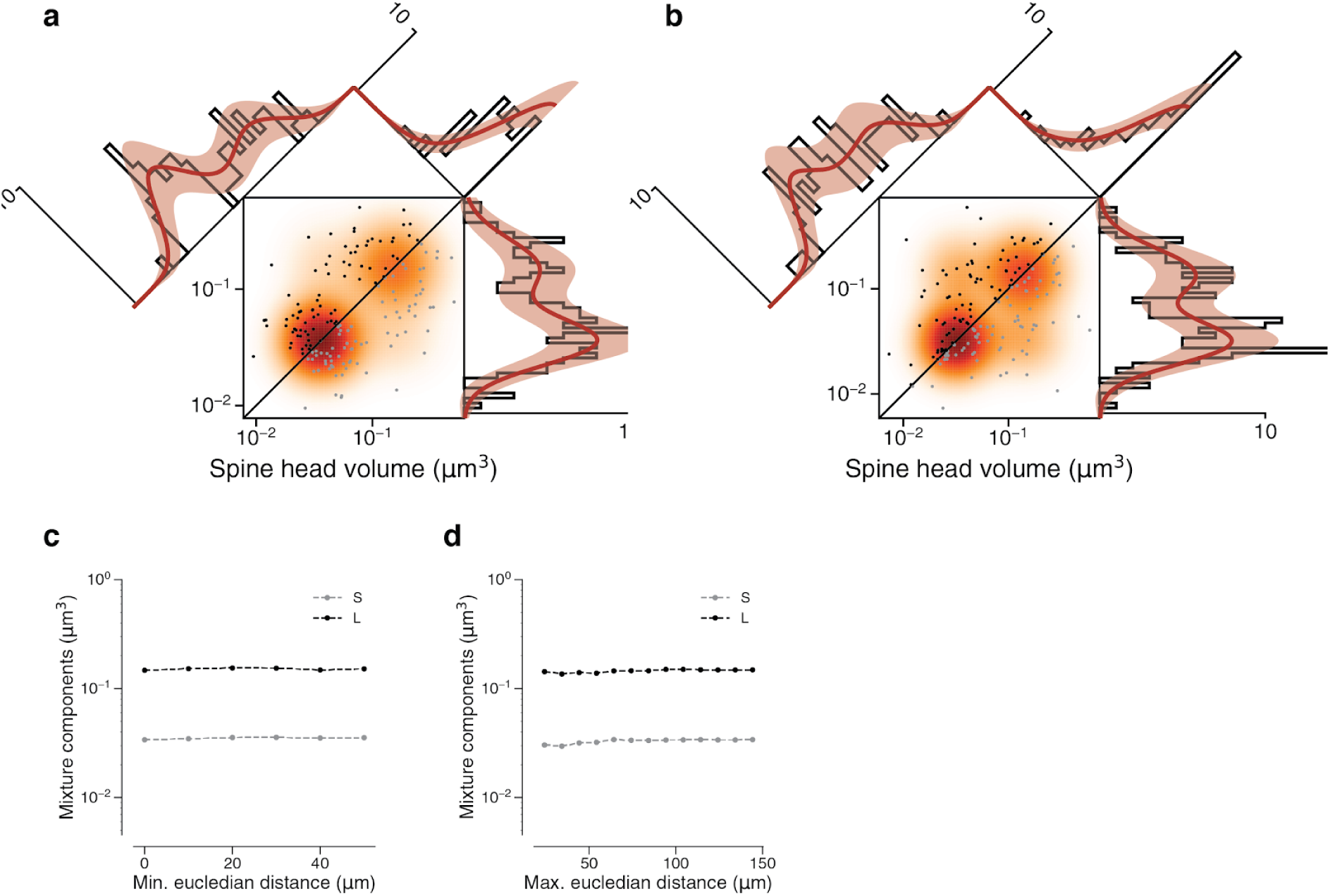
Synapses in a dual connection: near vs. far pairs. Spine volumes (left) and cleft sizes (right) **(a)** Dual connections of synapse pairs less than 46.5 μm apart (phi=0.534). **(b)** Dual connections of synapse pairs more than 46.5 μm apart (phi=0.745). **(c)** Mixture component means of model fits as a function of the minimum distance separating synapse pairs. **(d)** Mixture component means of model fits as a function of the maximum distance separating synapse pairs Error bars are ± 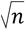 of the model fit.

**Extended Data Figure 9.**
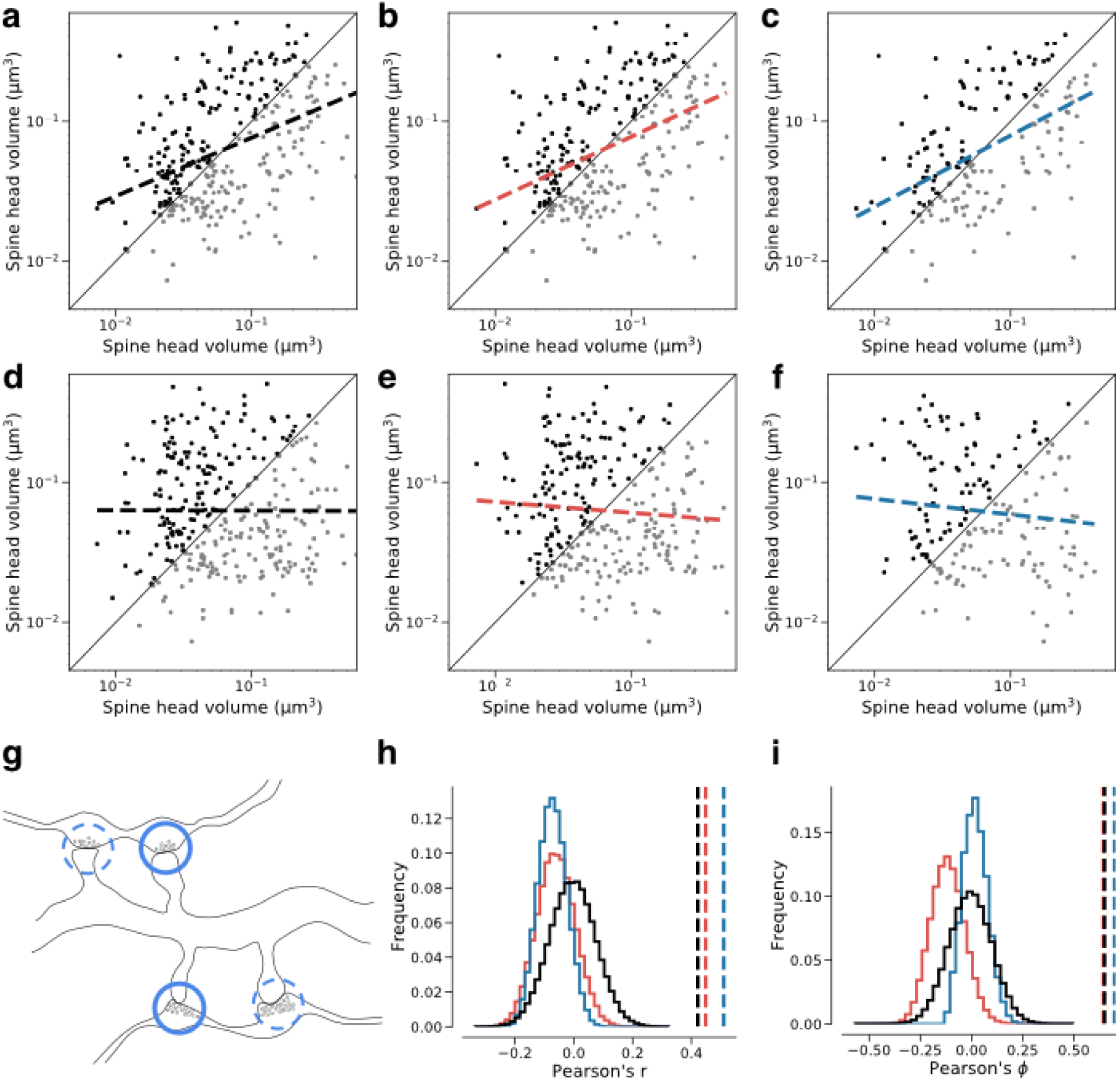
Dual connection correlations are not a result of axon or dendrite biases. Synapses are shuffled between dual connections to measure the correlations between synapses that have the same axon (red) or the same dendrite (blue) against a fully random baseline (black). Shuffled synapses are not allowed to be paired with other synapses within their original connection. Each shuffling procedure was performed on a subset of data where at least one valid shuffling exists for each synapse (e.g. where each dendrite receives at least two dual connections). (a-c) Joint distributions of spine head volumes in the dual connections used for shuffling. Slope of linear fit shows Pearson’s r value. **(a)** Subset of data used for random shuffling (all 160 synapse pairs). r= 0.42. **(b)** Subset of data used for axon-preserved shuffling (141 pairs). r= 0.45. **(c)** Subset of data used for dendrite-preserved shuffling (89 pairs). r= 0.51. **(d)** Example shuffle of data in (a). r= 0.00. **(e)** Example shuffle of data in (b). r= -0.08. **(f)** Example shuffle of data in (c). r= -0.11. **(g)** Diagram of a possible shuffle of two dual synaptic connections onto the same dendrite. **(h)** Distribution of Pearson’s r correlation for paired spine head volumes after shuffling (100,000 shuffles each). Dashed lines indicate correlation value for the unshuffled subset. **(i)** Distribution of Pearson’s phi correlation for paired spine head volumes after shuffling

**Extended Data Figure 10.**
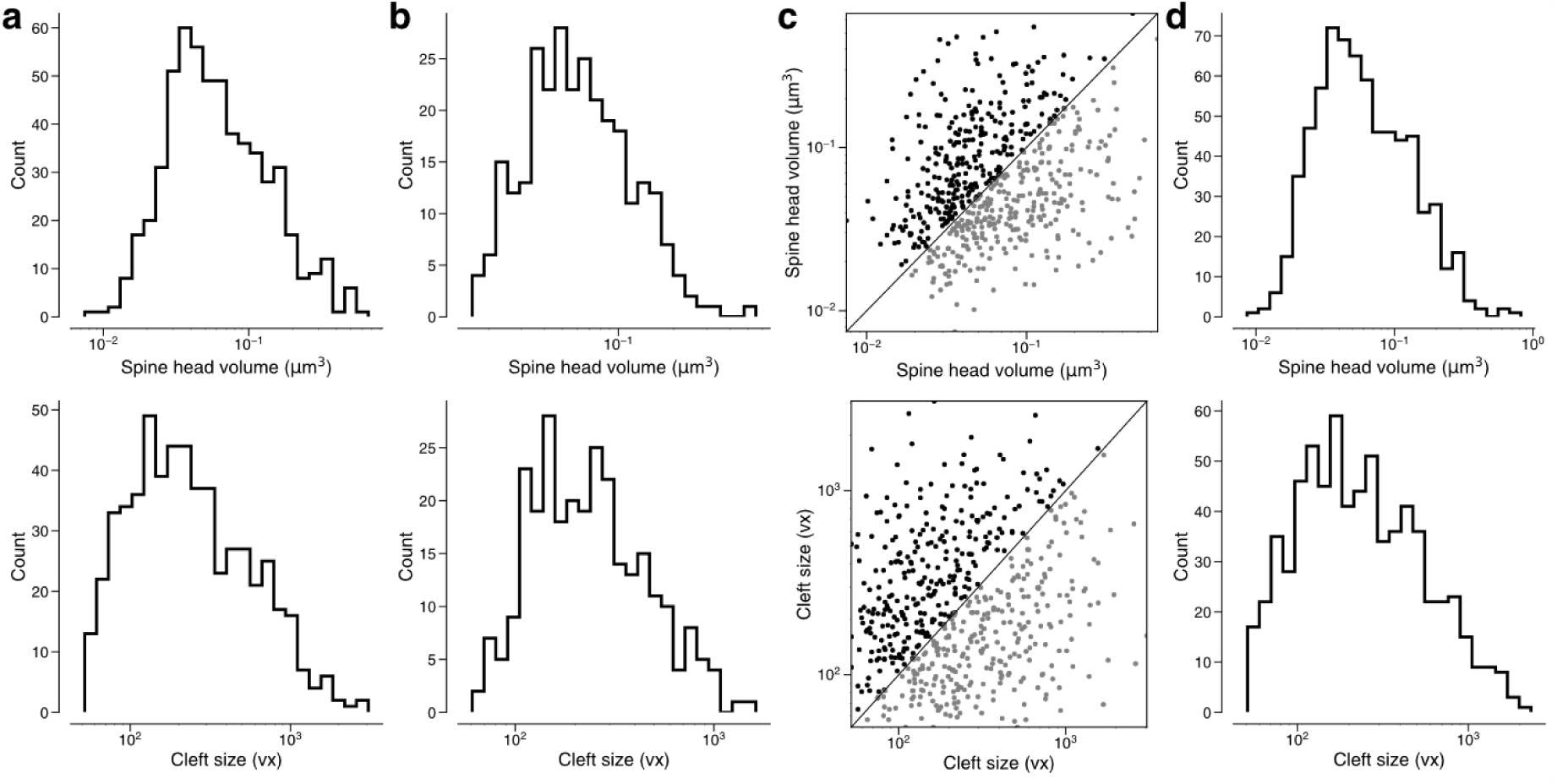
Removing constraints on the synaptic population eliminates bimodality and reduces correlations. Spine volumes (top) and cleft sizes (bottom) **(a)** Distribution of synapse sizes in dual connections received by L2/3 pyramidal cells, including those from orphan axons (566 synapses). **(b)** Distribution of geometric means of synapse sizes in same dual connections as in (**a**) (283 pairs). **(c)** Joint distribution of synapse sizes in dual connections received by L2/3 pyramidal cells, including those from orphan axons (283 pairs). **(d)** Distribution of synapse sizes for excitatory synapses received by L2/3 pyramidal cells, including those from orphan axons (700 synapses).

**Supplementary Information 1.**
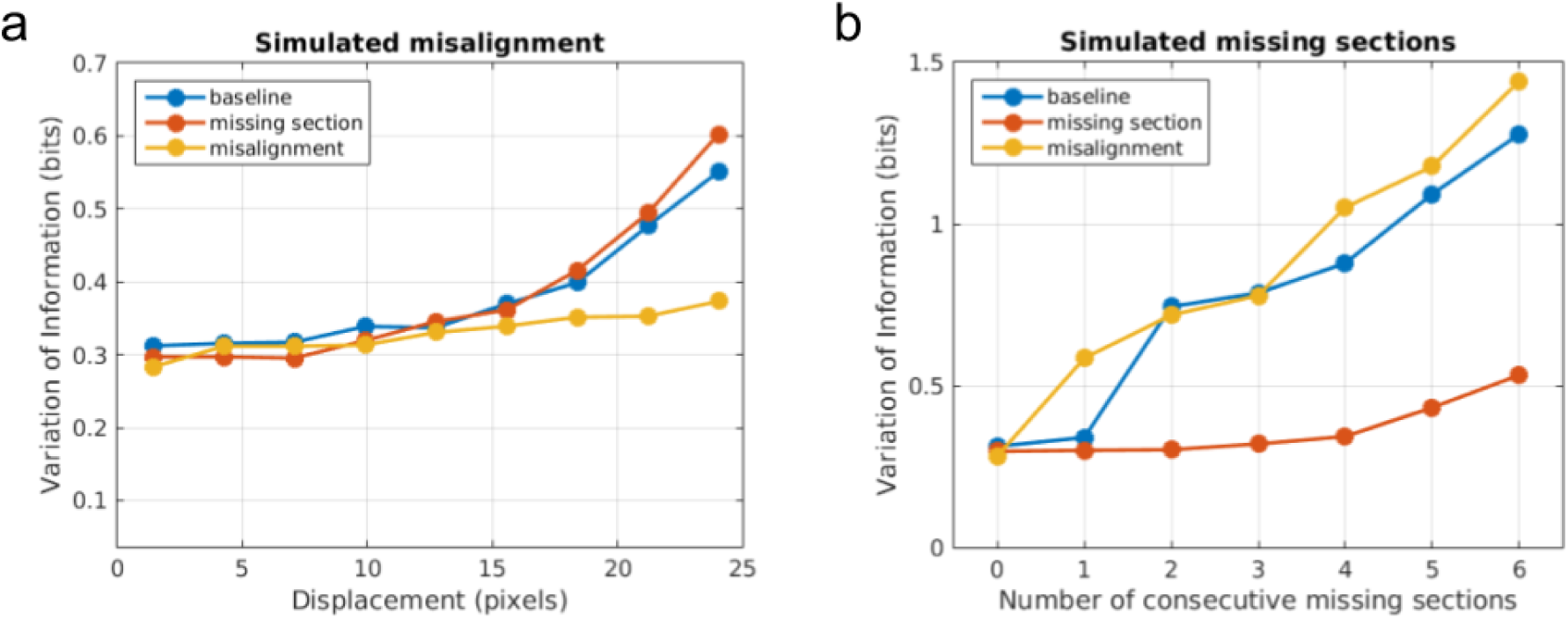
Quantitative evidence for the effectiveness of training data augmentation. Robustness of three boundary detectors trained with no data augmentation (“baseline”, blue), simulated missing section (“missing section”, red), and simulated misalignment (“misalignment”, yellow) to (a) increasingly large displacement of simulated misalignment and (b) increasing number of simulated consecutive missing sections.

**Methods.**

We performed preliminary experiments on the effect of training data augmentation by simulating image defects on the publicly available SNEMI3D challenge dataset (http://brainiac2.mit.edu/SNEMI3D). We partitioned the SNEMI3D training volume of 1024 x 1024 x 100 voxels into the center crop of 512 x 512 x 100 voxels for validation, and the rest for training. Then we trained three convolutional nets to detect neuronal boundaries, one without any data augmentation (“baseline”), and the other two with simulated missing section (“missing section”) and simulated misalignment (“misalignment”) data augmentation, respectively. After training the three nets, we measured the robustness of each net to varying degrees of simulated image defects on the validation set. In the first measurement, we simulated a misalignment at the middle of the validation volume with varying numbers of pixel displacement. In the second measurement, we introduced varying numbers of consecutive missing sections at the middle of the validation volume. For each configuration of simulation, we ran inference pipeline with the three nets to produce respective segmentations, and computed the Variation of Information error metric to measure the quality of the segmentations. For the measurement against simulated misalignment, we applied connected components to recompute the ground truth segmentation after introducing a misalignment, such that we separated a single object into two distinct objects if the object is completely broken by the misalignment (e.g. the displacement of misalignment larger than the diameter of neurite).

**Supplementary Information 2.**
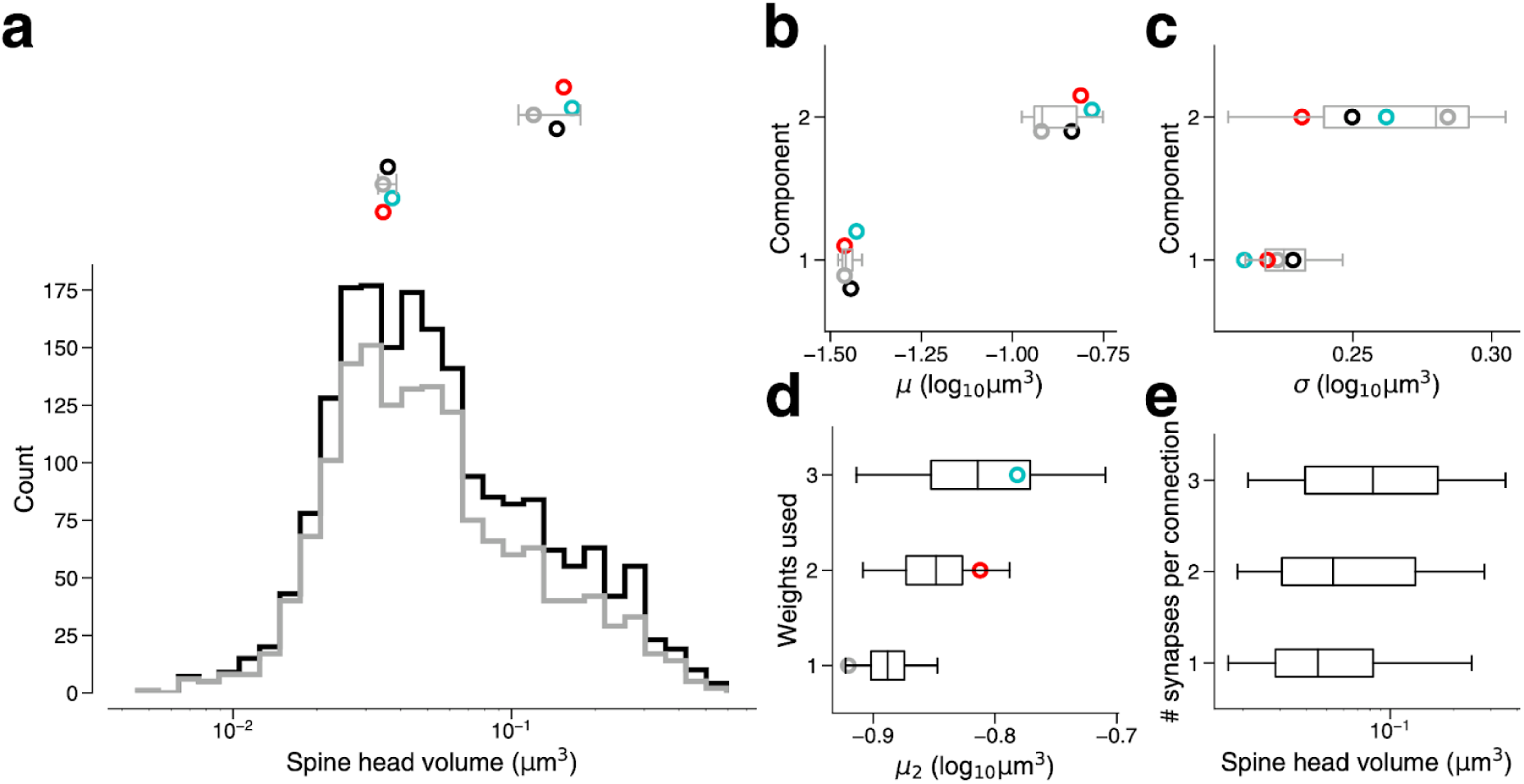
Synapse size by connection type. **(a)** Bottom: Spine head volume distribution for single connections (gray) along with synapses for all connections (black). Top: parameter estimates for the component means of single connections, as well as dual connections (red), and triple connections (cyan) and all connections (black). Gray line indicates 90% bootstrap interval over the single connection synapses (1000 samples). Points jittered for clarity. **(b)** Parameter estimates for component means of the same populations in (a). Box indicates interquartile range across bootstrap samples. Whiskers show 90% bootstrap interval. **(c)** Parameter estimates for component standard deviations of the same populations in (a). **(d)** Second component mean estimate from GMM fits on samples of the full dataset model. Full dataset model used for sampling had component weights taken from single connection model, dual connection model, or triple connection model. Points show parameter estimate from the original GMM fit for that connection type. **(e)** Spine head volume by the number of synapses in a connection.

**Supplementary Information 3.**
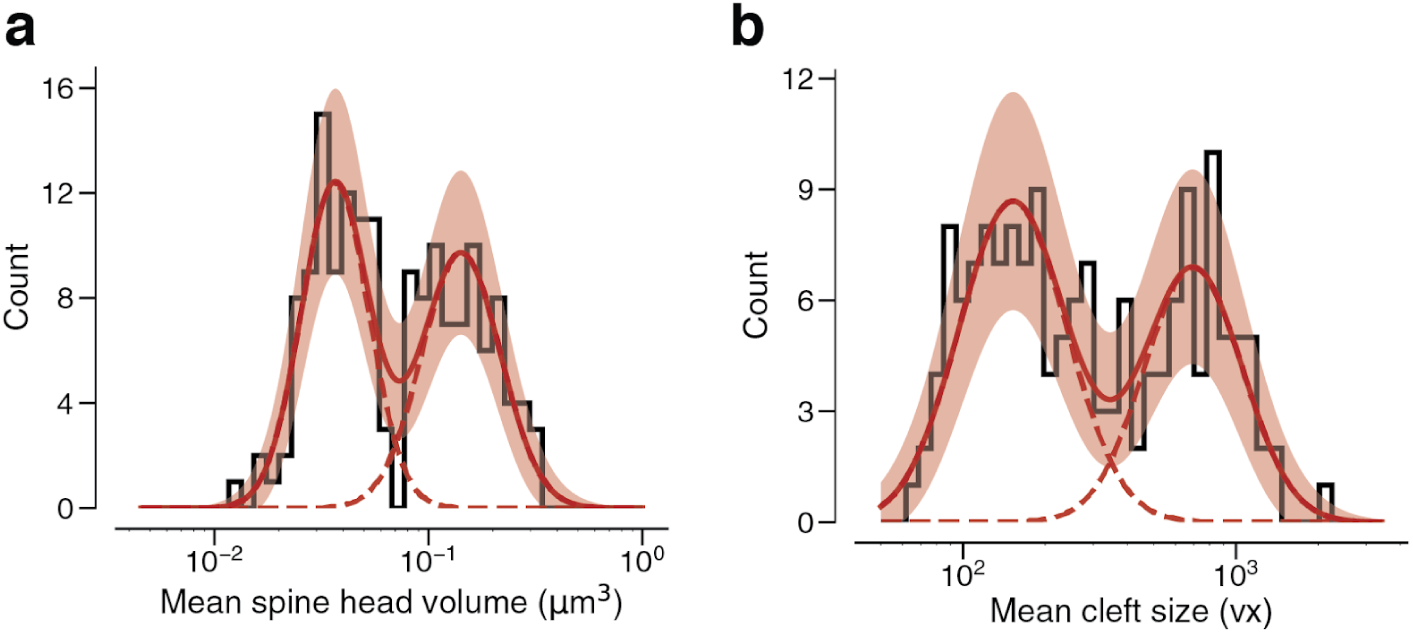
Arithmetic means. **(a)** Dual connections between L2/3 PyCs, each summarized by the arithmetic mean of two spine volumes, modeled by a mixture (red, solid) of two normal distributions (red, dashed). **(b)** Dual connections between L2/3 PyCs, each summarized by the arithmetic mean of two cleft sizes, modeled by a mixture (red, solid) of two normal distributions (red, dashed). Error bars are ± 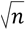 of the model fit.

**Supplementary Information 4.**
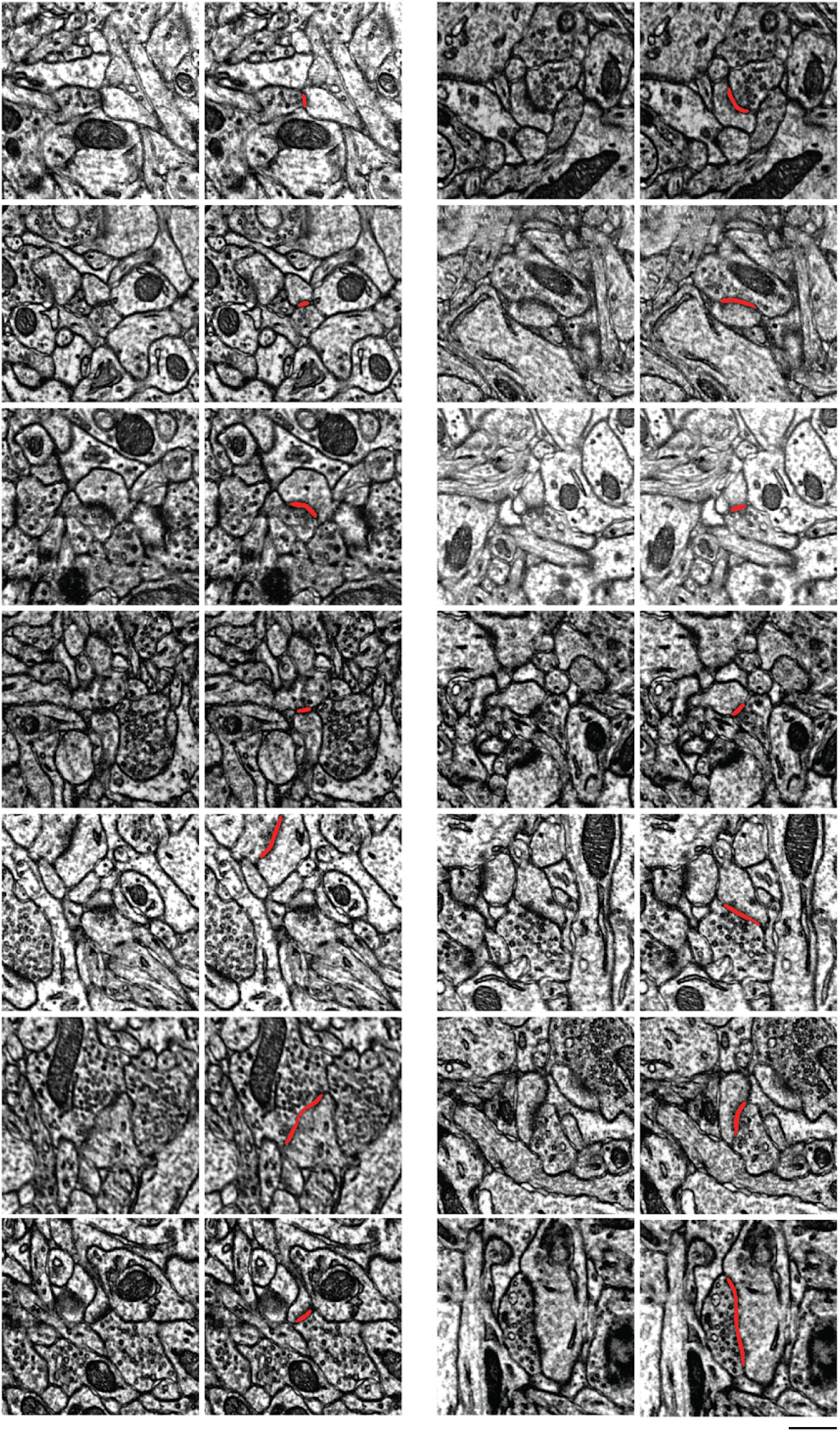
Examples of synapses between L2/3 pyramidal cells. In each pair of images, the left shows one section through a synapse, and the right adds the automatically detected cleft as an overlay. Note here that the clefts are associated with postsynaptic densities.

**Supplementary Information 5.**
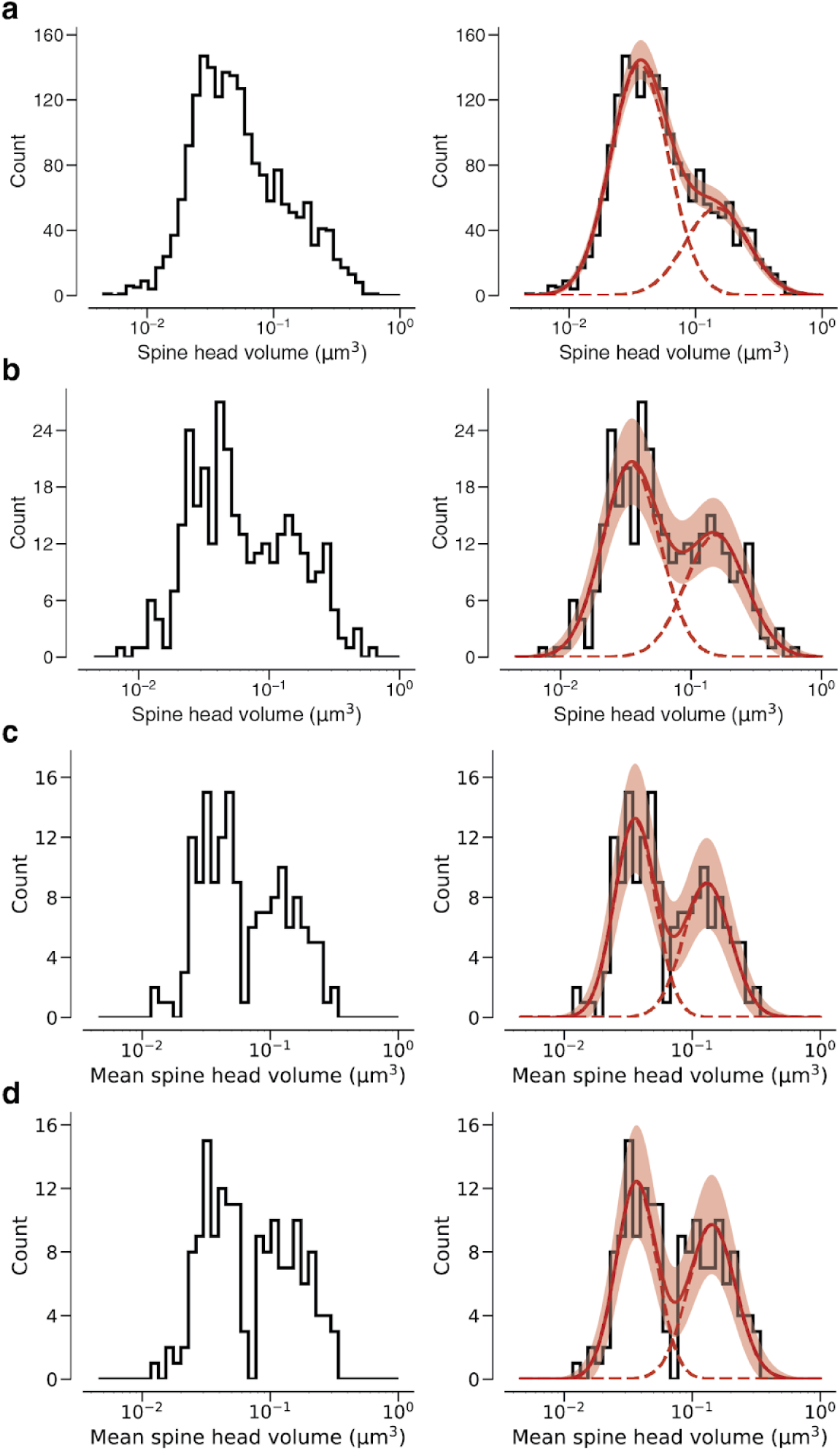
Fits vs raw data histograms. Plots are analogous to Fig. 3 and SI 3. **(a)** Histogram of same spine volumes, logarithmic scale. A mixture (red, solid) of two log-normal distributions (red, dashed) is shown. **(b)** Spine volumes belonging to dual connections between L2/3 PyCs, modeled by a mixture (red, solid) of two log-normal distributions (red, dashed). **(c)** Dual connections between L2/3 PyCs, each summarized by the geometric mean of two spine volumes, modeled by a mixture (red, solid) of two log-normal distributions (red, dashed). **(d)** Dual connections between L2/3 PyCs, each summarized by the arithmetic mean of two spine volumes, modeled by a mixture (red, solid) of two log-normal distributions (red, dashed).

**Supplementary Information 6.**
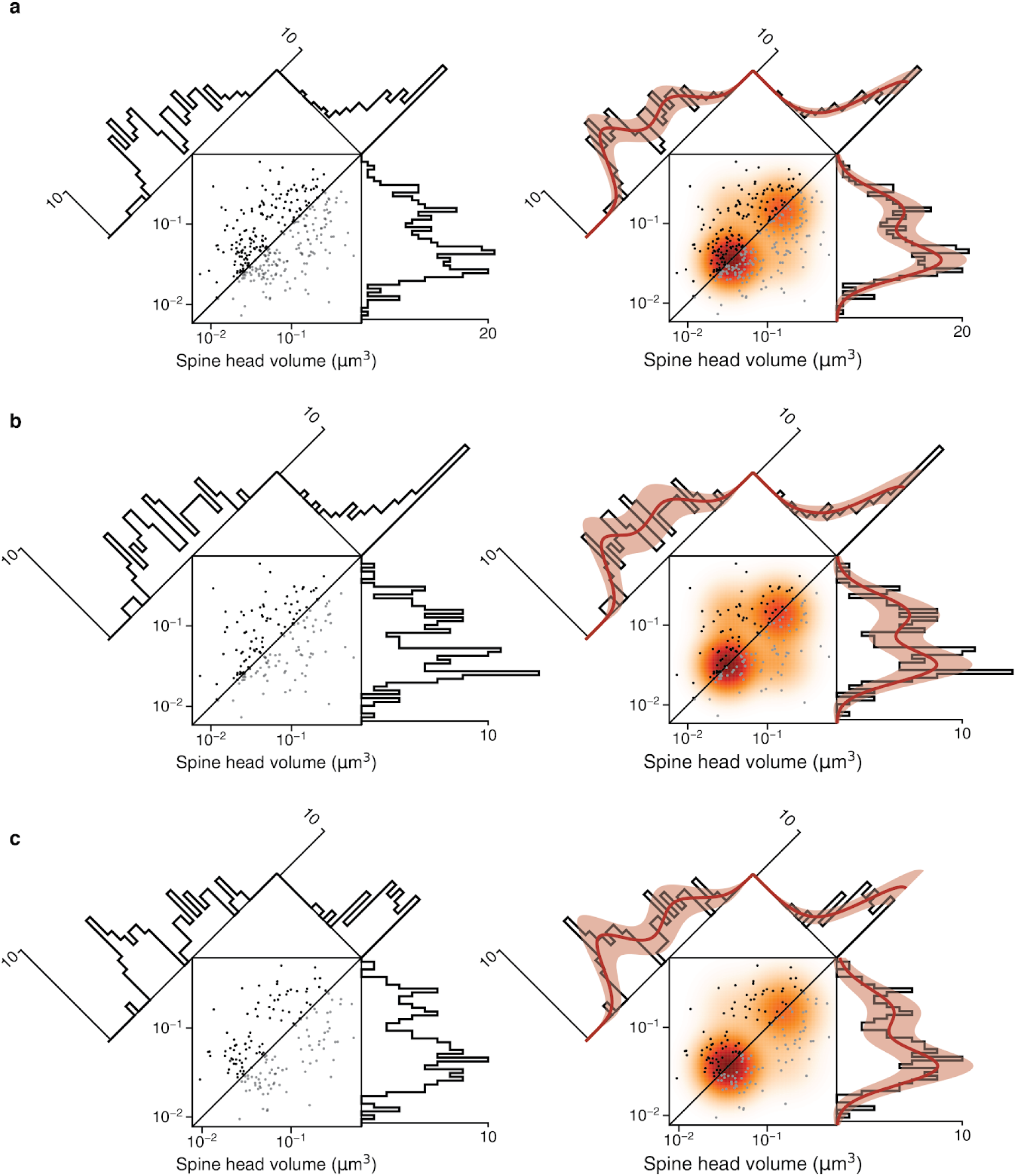
Fits vs raw distributions. Plots are analogous to Fig. 4 and Ext. Data Fig. 8. **(a)** Synapse pairs from all dual connections. **(b)** Dual connections of synapse pairs less than the median distance apart. **(c)** Dual connections of synapse pairs more than the median distance apart

**Supplementary Information 7.**
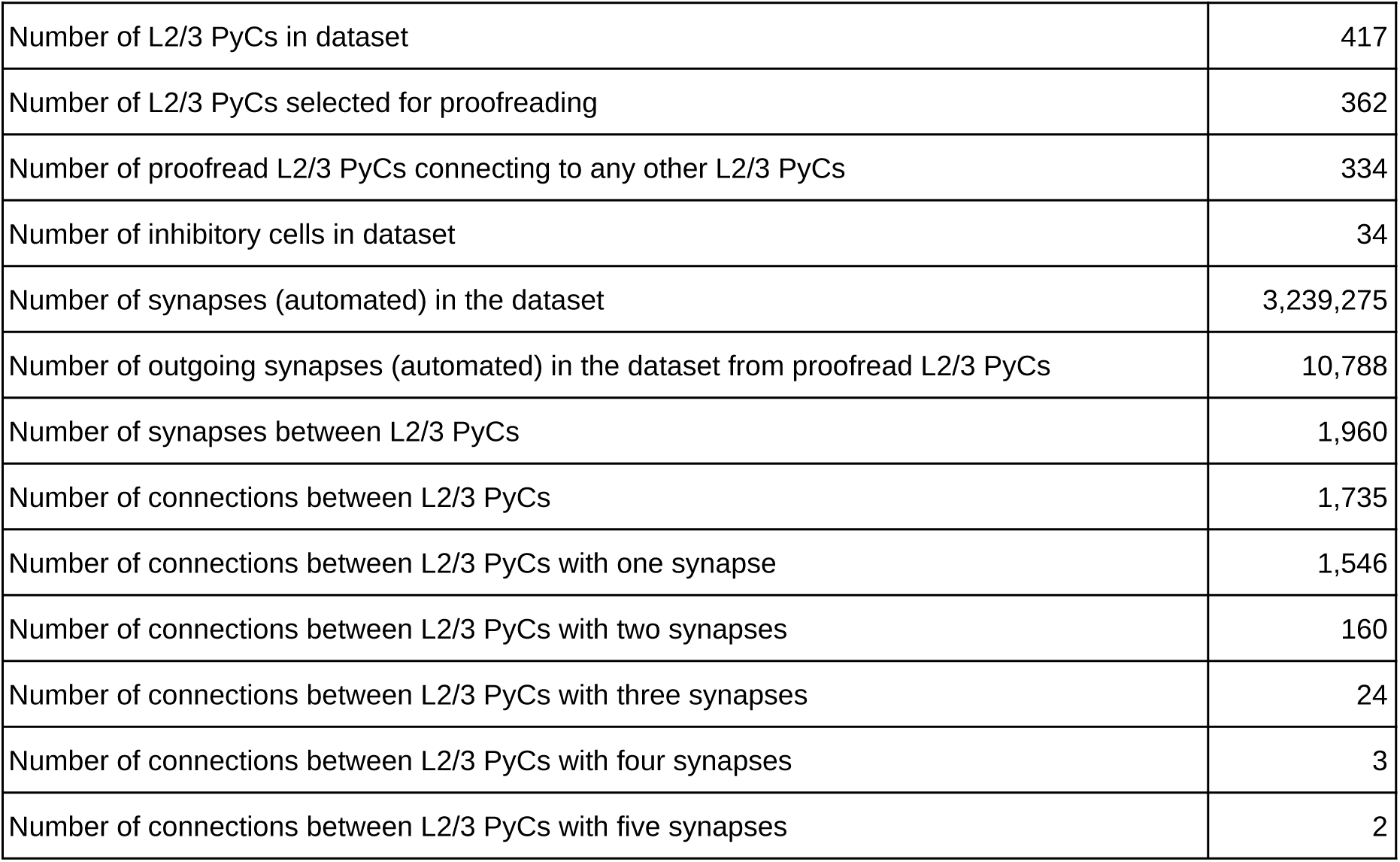
Overview of number of data points obtained in this study.

|  |  |
| --- | --- |
| Number of L2/3 PyCs in dataset | 417 |
| Number of L2/3 PyCs selected for proofreading | 362 |
| Number of proofread L2/3 PyCs connecting to any other L2/3 PyCs | 334 |
| Number of inhibitory cells in dataset | 34 |
| Number of synapses (automated) in the dataset | 3,239,275 |
| Number of outgoing synapses (automated) in the dataset from proofread L2/3 PyCs | 10,788 |
| Number of synapses between L2/3 PyCs | 1,960 |
| Number of connections between L2/3 PyCs | 1,735 |
| Number of connections between L2/3 PyCs with one synapse | 1,546 |
| Number of connections between L2/3 PyCs with two synapses | 160 |
| Number of connections between L2/3 PyCs with three synapses | 24 |
| Number of connections between L2/3 PyCs with four synapses | 3 |
| Number of connections between L2/3 PyCs with five synapses | 2 |

**Supplementary Information 8.**
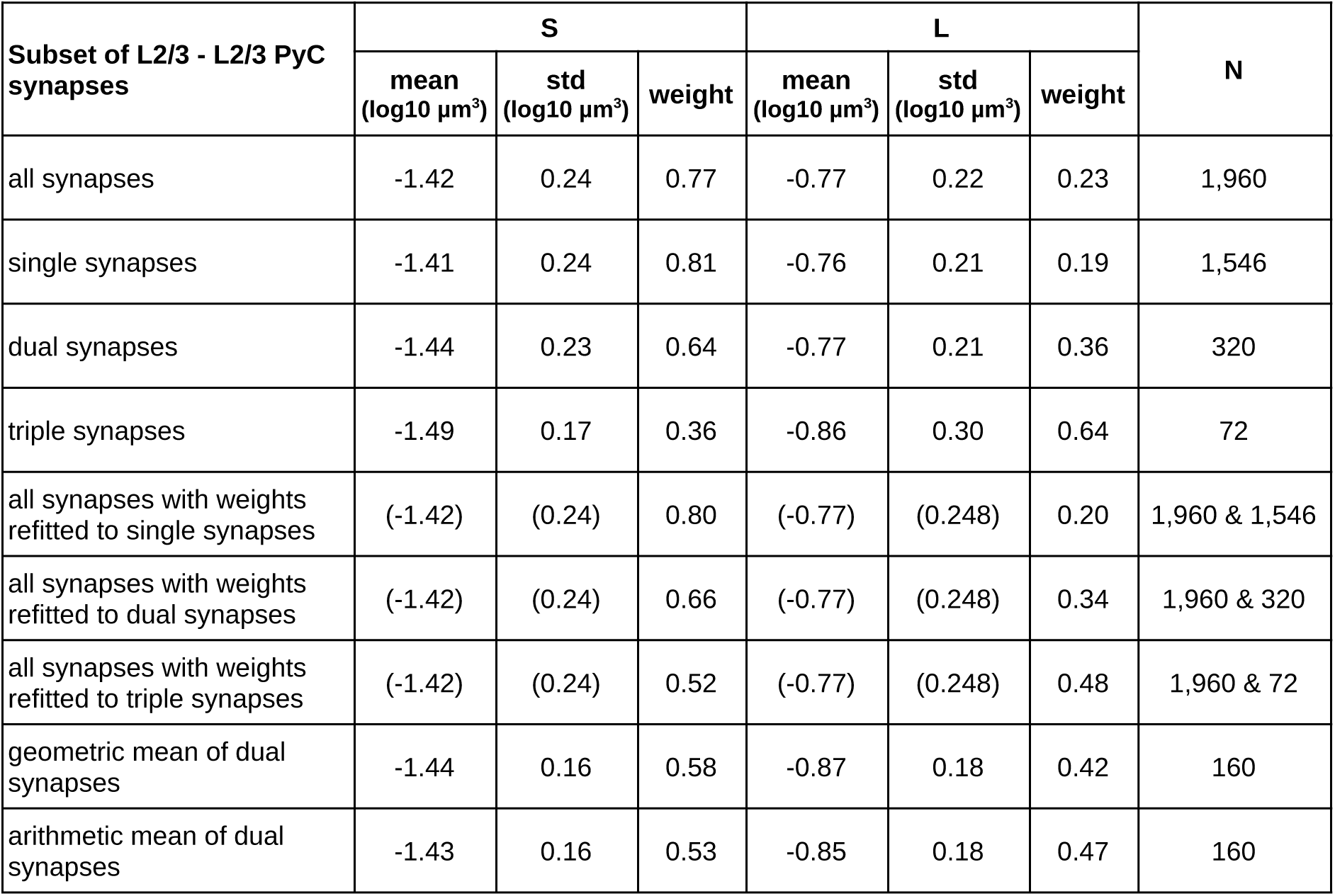
Overview of results from lognormal mixture fits for different synapse subpopulations.

**Supplementary Information 9.** Overview of results from HMM lognormal component fits for different dual synaptic connection subpopulations.

**Supplementary Information 8. Overview of results from lognormal mixture fits for different synapse subpopulations**
| Subset of L2/3 - L2/3<br>PyC dual synaptic<br>connections | S |  | L |  | weights |  |  | Pearson<br>-Phi | N |
| --- | --- | --- | --- | --- | --- | --- | --- | --- | --- |
| | mean<br>(log10 $\mu\text{m}^3$) | std<br>(log10 $\mu\text{m}^3$) | mean<br>(log10 $\mu\text{m}^3$) | std<br>(log10 $\mu\text{m}^3$) | SS | SL+<br>LS | LL | | |
| all connections | -1.470 | 0.216 | -0.833 | 0.244 | 0.490 | 0.177 | 0.333 | 0.637 | 160 |
| dist < median dist | -1.506 | 0.212 | -0.861 | 0.243 | 0.427 | 0.232 | 0.342 | 0.534 | 80 |
| dist > median dist | -1.449 | 0.207 | -0.818 | 0.251 | 0.529 | 0.123 | 0.348 | 0.745 | 80 |

## Footnotes

1 These numbers should be taken with the caveat that the observed number of synapses for a connection is a lower bound for the true number of synapses, because two PyCs with cell bodies in our EM volume could synapse with each other outside the bounds of the volume.

2 Spine head volume excludes the spine neck, which is at most only weakly correlated in size with other synaptic structures (Arellano et al. 2007).

3 For example, if the two normal distributions have the same weight and standard deviation, then the mixture is unimodal if and only if the separation between the means is twice the standard deviation.

4 This relationship was found for the *observed* number of synapses. On average, this number is expected to increase with the *true* number of synapses. Therefore mean spine volume is also expected to increase with the true number of synapses per connection.

5 The *x*=*y* and vertical histograms look bimodal because they are different projections of the same two “bumps” in the joint distribution. If the probability of the mixed state (LS/SL) were high, there would be two additional off-diagonal bumps in the joint distribution, and the *x*=*y* diagonal histogram would acquire another peak in the middle. In reality the probability of the mixed state is low, so the *x*=*y* diagonal histogram is well-modeled by two mixture components. The widths of the bumps are the same in both projections, but the distance between the bumps is longer in the *x*=*y* diagonal histogram by a factor of root two. This explains why the mixture components are better separated in the distribution of geometric means (Fig. 3e, 4e) than in the marginal distribution (Fig. 3d, 4d), and hence why the statistical significance of bimodality is stronger for the geometric means.

